# Structural and functional insights into the role of Cysteine-Rich Receptor-Like Kinase 18 (CRK18) in Arabidopsis

**DOI:** 10.64898/2026.07.30.741491

**Authors:** Jente Stouthamer, Judith Lanooij, Gonzalo Vílchez-Pinto, Rahul S. Lathe, Jan Eric Maika, Dušan Živković, Michelle von Arx, Sjef Boeren, Mark Roosjen, Anniek Oosterwijk, Simon Lindhoud, Corinne Geertsema, Ran Lu, Che-Yang Liao, Monique van Oers, Rüdiger Simon, Jani R. Bolla, Antonio Molina, Julien Gronnier, Jose Lozano Torres, María Garrido-Arandia, Elwira Smakowska-Luzan

## Abstract

Plants perceive and integrate diverse environmental signals through receptor kinase (RK) networks at the plasma membrane. Within this, Cysteine-Rich Receptor-Like Kinases (CRKs) constitute a large but not well-understood subfamily characterised by extracellular domains (ECDs) enriched in cysteine residues. CRKs have been implicated in plant responses to biotic and abiotic stress, as well as in developmental processes. Additionally, several CRKs have been proposed to act as redox sensors. Here, we investigate the homodimerization mechanism of Arabidopsis CRK18 and its regulation by redox conditions. By modulating pH and redox state, we assessed the stability and binding dynamics of the CRK18 ECD and its cysteine mutants. We also tested the plasma membrane localisation of all the cysteine mutants involved in the predicted disulfide bonds, and only CRK18C227A, C228A-ECD resembled the plasma membrane localization of wild-type CRK18. We combine co-immunoprecipitation, Förster resonance energy transfer–fluorescence lifetime imaging microscopy and microscale thermophoresis to quantify CRK18 self-association in planta and in vitro. Furthermore, we place CRK18 dimerization in a broader signalling context by identifying CRK18 interaction partners and CRK18-dependent signalling outputs. Using Arabidopsis CRK18 overexpression lines, we perform (phospho)proteomic and immunoprecipitation–mass spectrometry (IP-MS) analyses to map CRK18-centred signalling networks associated with stress-related responses. The constitutive activation of the CRK18 kinase domain and its interaction with many putative (cell wall) glycan-sensing RKs, including PERK15, suggest a regulatory role for CRK18. The absence of significant changes in the proteome and phosphoproteome of CRK18 overexpression lines in the absence of any trigger, and its restricted mobility after elicitation, suggest that CRK18 requires a stimulus for activation and possibly induces membrane microdomain reorganisation. This is in line with the infection assay with the nematode Heterodera schachtii, which causes modification and targeted damage to plant cell walls during infection, revealing that CRK18 acts as negative regulator of this process.

## Introduction

Plants continuously monitor their environment, integrating information about light, temperature, water, and nutrient availability, as well as the presence of symbiotic or pathogenic organisms(Heil and Ton 2008; Couto and Zipfel 2016; Tarkowski et al. 2022). They not only sense multiple cues simultaneously but also mount dynamic and highly specific responses. At the molecular level, these responses are mediated by intricate signalling networks, in which plasma membrane-localized receptor kinases (RKs) play a central role by perceiving extracellular signalling molecules(Han et al. 2014; Brandt and Hothorn 2016; Hohmann et al. 2017; Wolf 2017; Smakowska-Luzan et al. 2018; Mittler et al. 2022). RKs recognise a broad range of ligands, including hormones, small peptides and sugars, via their extracellular domains (ECDs). Ligand binding induces the formation of receptor complexes with co-receptors and additional regulatory RKs, leading to transphosphorylation of intracellular kinase domains(Han et al. 2014). Activated complexes initiate downstream signalling cascades that regulate diverse processes such as Reactive Oxygen Species (ROS) production, apoplastic pH changes, cytosolic Ca²⁺ burst, stomatal movements and transcriptional responses. Through these mechanisms, RK complexes translate environmental information into appropriate cellular and physiological outputs(Pok et al. 2026).

Among plant RKs, Cysteine-Rich Receptor-Like Kinases (CRKs) represent a large but still poorly understood subfamily(Zhang et al.; Chen 2001; Martin-Ramirez et al. 2025). The Arabidopsis CRK family comprises 44 members. When first described, CRK ECDs were defined by the presence of two Domain of Unknown Function 26 (DUF26) modules, each enriched in cysteine residues and containing a characteristic C–X₈–C–X₂–C motif, from which the family derives its name(Stouthamer et al. 2026). Since their discovery, CRKs have been implicated in both stress responses and developmental processes(Zhang et al.; Chen 2001). Recent Redox Interactome Assay (RIA) studies have demonstrated that CRKs interact with each other in a ROS-dependent manner and that ROS rewires the CRK interaction network. Also, several CRKs were identified as putative ROS sensors, with CRK28 playing a redox-dependent role in senescence and autoimmune responses(Martin-Ramirez et al. 2026).

Previous studies have shown that CRK gene expression is induced by biotic and abiotic stimuli, including pathogens, pathogen-associated molecular patterns (PAMPs), ozone, and phytohormones(Wrzaczek et al. 2010; Bourdais et al. 2015). Genetic analyses have linked individual CRKs to plant immunity: loss-of-function mutants exhibit altered ROS production upon PAMP treatment and altered susceptibility to bacterial and fungal pathogens, whereas overexpression of specific CRKs can enhance resistance or trigger cell death, likely in an expression-level–dependent manner(Acharya et al. 2007; Ederli et al. 2011; Lee et al. 2017; Pelagio-Flores et al. 2020; Zhao et al. 2022; Piovesana et al. 2023). Several CRKs associate with pattern-recognition receptors and co-receptors, such as FLS2, BAK1, and BIK1(Lee et al. 2017; Yadeta et al. 2017a). Furthermore, CRK2 has been shown to phosphorylate the NADPH oxidase RBOHD and to modulate PAMP-induced ROS production (Kimura et al. 2020). CRKs have also been associated with abiotic stress responses, including salt and drought tolerance, and with developmental processes, such as biomass accumulation, senescence, and root architecture(Idänheimo et al. 2014; Lu et al. 2016; Pelagio-Flores et al. 2020; Zhou et al. 2023; Imai et al. 2024). Together, these observations support the view that CRKs act as context-dependent regulators within RK signalling networks, although their precise biochemical functions and signalling pathways remain elusive.

A central focus of our work was the homodimerisation mechanism of CRKs, using CRK18 as a model for study. CRK18 is an attractive candidate because its ECD is the most negatively charged among the CRKs and contains cysteine residues in both conserved and non-conserved motifs. AlphaFold-based structural predictions, multiscale molecular dynamics simulations, and RIA revealed that CRK18 forms a homodimeric complex at the ECD level and that this complex formation is disrupted in the presence of ROS. We combined co-immunoprecipitation, Förster resonance energy transfer–fluorescence lifetime imaging microscopy (FRET–FLIM) and microscale thermophoresis (MST) to assess CRK18 self-association in planta and *in vitro.* We examined how pH, redox conditions, and mutations of vicinal cysteines affect dimer formation, thereby elucidating the principles of CRK18 dimerization and its potential for redox sensitivity. These experiments revealed how specific cysteine residues and disulphide bonds contribute to CRK18 stability and dynamics. Finally, using (phospho)proteomic and immunoprecipitation–mass spectrometry analyses we placed CRK18 in a broader signalling context by identifying CRK18 interaction partners and thereby identifying putative roles in cellular and physiological responses. We uncovered that CRK18, possibly because of its association with a PERK family member, serves as a negative regulator of *H. schachtii* infection, possibly contributing to monitoring cell wall integrity. Together, our results provide mechanistic insight into the dimerization behaviour of CRK18, clarify the structural contributions of its cysteines and disulphide bonds, and establish a framework for understanding how CRK18 and potentially other RKs integrate complex formation into plant stress-related signalling pathways.

## Materials and methods

### Plasmid construction and baculovirus stock preparation

To express CRK18 in *N. benthamiana*, CRK18 genomic DNA from Arabidopsis was amplified with overhangs for modified pGIIB expression vectors (de Rybel et al. 2011). ER-Marker gene AtVMA21a was obtained from J.W. Borst. The construct was cloned into the vector with C-terminal 3xFLAG-mScarlet-I or mNeonGreen-V5 tags using HiFi DNA assembly according to the manufacturer’s instructions (NEB). To introduce Cysteine (C) to Alanine (A) mutations or replace the conserved Lysine (K) of the kinase ATP-binding site with Aspartic acid (D), CRK DNA fragments were amplified with primers containing the respective point mutations. Fragments were then reassembled using HiFi (NEB).

For FRET-FLIM experiments, CRK18 cDNA was amplified from Arabidopsis and cloned into estradiol inducible expression vectors (Bleckmann et al. 2009) with C-terminal fluorescent protein tags (GFP, mCherry or tandem GFP-mCherry) using Gateway as described by manufacturer’s instructions (Invitrogen).

To generate expression vectors for baculovirus stocks, the CRK18-ECD DNA (encoding residues 26-284) was amplified from cDNA of *Arabidopsis thaliana*. cDNA was amplified with extensions for sequences encoding for an N-terminal Melittin signalling peptide and a C-terminal 9xHis tag and attB overhang sites. Mel-CRK18-ECD-9xHis was inserted into the pDONR221 entry vector and subsequently into the pOET1 vector (Oxford Expression Systems) using the Gateway method (Invitrogen). Cys to Ala point mutations were introduced using the site-directed mutagenesis method.

Baculovirus stocks were generated using the flashBAC Ultra system (Oxford Expression Systems) in *Spodoptera frugiperda* Sf9 insect (Expression Systems) cells according to manufacturers’ instructions. Viral titers were determined using Sf9-EasyTiter cells(Hopkins and Esposito 2009).

### Protein expression and purification

*Trichoplusia ni* High Five Cells (BTI-TN-5B1-4) were cultured in ESF291 Insect Cell Culture Medium (Expression Systems) at 27°C in suspension culture. Cells were infected with viral stocks to a multiplicity of infection of 10 and grown for 72 h at 27 °C in suspension culture. Next, the protein-containing supernatant was recovered by centrifugation at 1500g for 30 min. Supernatant conditions were adjusted by addition of NaCl to 300 mM, DTT to 1 mM, and Imidazole to 10 mM. The supernatant was incubated with 3 mL/L Ni Sepharose Excel resin (Cytiva Life Sciences) for 2h at 4 °C while shaking. The resin was transferred to a column and washed with wash buffer (20 mM Tris, 500 mM NaCl, 10 mM imidazole, 1 mM DTT, pH 8). Protein was eluted with Elution buffer (20 mM Tris, 500 mM NaCl, 350 mM Imidazole, 1 mM DTT, pH 8). The eluted protein was concentrated using 10-kDa Amicon centrifuge filters (Millipore) and further purified using an AKTA Pure FPLC system on a Superdex 200 preparative-grade column (Cytiva Life Sciences) in a buffer containing 20 mM HEPES, 150 mM NaCl, pH 7.5. Collected fractions were assessed with SDS-PAGE and pure fractions were pooled and concentrated.

### nanoDSF

To assess the stability of the purified CRK-ECDs, nanoDSF were performed. Protein samples in either 20 mM HEPES, 150 mM NaCl, 0.1% Pluronic acid, pH 7.5 or 20 mM NaOAc, 150 mM NaCl, 0.1% Pluronic acid, pH 5 were prepared at a concentration of 0.4 mg/mL protein. Samples were measured in triplicate in standard capillaries using a Prometheus Panta device (NanoTemper). Temperature was increased from 20-95 °C at a ramp rate of 0.5 °C/min. Samples were excited at 280 nm and fluorescence emission was measured at 330 nm and 350 nm. Data was analysed using Panta analysis software (NanoTemper). The T_m_ was derived from the inflexion point in the sigmoidal curve by determining the min/max of the first derivative of the fluorescence intensity at 350nm, 330nm and the ratio thereof.

### Microscale Thermophoresis

To measure CRK18-ECD interactions, microscale thermophoresis (MST) was used. CRK18-ECD and CRK18^C227A,C228A^-ECD were labelled with DyLight650 NHS dye (Invitrogen) according to manufacturer’s instructions, and free dye was removed with zebra dye removal columns. To prepare samples for MST, a serial dilution of unlabelled protein was prepared, and labelled protein was added at a fixed concentration. For all measurements, the final concentration of labelled protein in the samples was 20 nM. The unlabelled protein starting concentration, depended on the measurement: CRK18-CRK18 = 100 µM, CRK18C227A,C228A = 86 µM, CRK18-18 pH5 = 72 µM, CRK18-18 + DTT = 88 µM. Single repeats of CRK18 with DTT or H_2_O_2_ treatments = 90 µM. Samples were prepared in either a 20 mM HEPES, 150 mM NaCl pH7.5 buffer or a 20 mM NaOAc, 150 mM NaCl buffer. 0.1% Pluoronic acid was added to prevent adhesion of the protein to the capillaries. Samples treated with 100 µM H_2_O_2_ or 0.5 mM DTT were incubated at RT for 1h before measuring. Samples were loaded into Monolith premium capillaries (NanoTemper) and MST measurements were performed using a Monolith NT.155 (NanoTemper). MST power was kept at 40%, excitation power varied per sample between 50-100%. Measurements were analysed by MO.Affinity Analysis v3.0.5 (NanoTemper), and further processed in Prism 10.4 (GraphPad).

### Biophysical characterisation of CRK18-ECD

SEC-MALS was used to determine protein molar mass, degree of glycosylation and oligomeric state. A CRK18-ECD sample at a concentration of 2.0 mg/mL in 20 mM HEPES, 150 mM NaCl, pH 7.5. was resolved on a Superdex 200 increase 10/300 GL column (Cytiva Life Sciences) using a 1260 infinity II high-pressure liquid chromatography system (Agilent). Particle diffraction and scattering was detected using an Optilab 1090 Differential refractive index detector and a miniDawn 1065 1065 Multi-Angle Light Scattering system (Wyatt Technologies). SEC-MALS measurements were performed in duplicate, and data were analysed using Astra 8.0 software (Wyatt Technologies). To determine the molar mass and glycosylation, dn/dc values of 0.185 ml/g for protein and 0.14 ml/g for glycans, and an extinction coefficient for CRK18-ECD at 280 nm of 0.945 ml/(mg/cm) were used.

Mass Photometry experiments were performed with CRK18-ECD at concentrations of 1.25 ug/mL and 12.5 ug/mL in NaOAc buffer pH 5. The sample was prepared as serial dilutions at room temperature, and it was analysed within 5 minutes upon dilution. MP was measured using a Refeyn Two^MP^ mass photometer and AcquireMP 2.3 software. Acquisition was done under normal size of field of view, with a 60 second recording. The video was processed with DiscoverMP 2.3 (Refeyn Ltd) (Frances Pugh et al. 2024), using default processing settings.

### Transient expression of CRKs in *N. benthamiana*

To transiently express CRKs in *N. benthamiana*, expression constructs were transformed into *A. tumefaciens* strain GV3101-pSoup. Cultures were grown for 72 hours at 28 °C. *Agrobacterium* solutions with a final OD of 0.3 for constructs and 0.5 for P19 were prepared in 10 mM MES, 10 mM MgCl, 100 µM acetosyringone. Agrobacterium solutions were incubated for 2 hours at room temperature. Leaves of 4-week-old *N. benthamiana* plants were infiltrated.

### Confocal microscopy

Leaf discs of four-week-old *N. benthamiana* plants 2 days post infection were measured using a Leica SP8 confocal microscope. mScI tagged recombinant protein (CRK18 and mutant variants) was excited at 561nm, and emission was detected between 580-630 nm using a hybrid detector. Recombinant ER-marker-mNG was excited with a 488 nm laser and emission was detected between 490-530 nm using a hybrid detector. A 63x water immersion objective (NA 1.2) was used. Images were processed in ImageJ. For each sample three images were recorded from three different plants.

### Co-Immunoprecipitation in *N. benthamiana*

Plants were infiltrated with Agrobacterium and the expression and localisation of 35S:CRK18^K367N^-3xFLAG-mScI, 35S:CRK18^C227A,C228A,K367N^-3xFLAG-mScI, and 35S:CRK18^K367N^-3xFLAG-mNG was checked by microscopy as explained above. After 2 days, infiltrated leaves were harvested, frozen in LN and ground into fine powder. Tissue powder was lysed for 1 h in lysis buffer (50 mM Tris pH7.5, 150 mM NaCl, 10% glycerol, 10 mM EDTA, 1 mM Sodium Molybdate, 20 mM NaF, 0.5% w/v PVPP, 10 mM DTT, 1% IGEPAL, protease inhibitor tablets (1/50 mL, Sigma). Plant material was filtered from the protein extract with miracloth. Anti-Flag agarose beads (Chromotek) were added to the protein extract and incubated for 3 hours at 4°C. Beads were washed four times in 20 mM Tris, 150 mM NaCl pH 7.5. To extract the protein, beads were incubated with SDS-PAGE sample buffer at 95°C for 10 min. Beads were removed by centrifugation. The protein samples were loaded on SDS-PAGE gels, and the protein was visualised by western blot followed by antibody illumination (anti FLAG-HRP and anti-V5-HRP, Invitrogen).

### FRET-FLIM in N. benthamiana

In order to measure interaction of CRK18 *in vivo* we employed FRET-FLIM. Transient expression in *N. benthamiana* of recombinant CRK18-GFP, CRK18-mCherry, CRK18-GFP-mCherry, and kinase-inactivated mutant variants with lysine 367 substituted by asparagine was performed as described in Blümke et al. (Blümke et al. 2021).

Lifetime measurements were performed using a Zeiss LSM 780 confocal microscope (40× water immersion objective, Zeiss C-PlanApo, NA 1.2). For Time-Corelated-Single-Photon-Counting (TCSPC) measurements a PicoQuant Hydra Harp 400 (PicoQuant, Berlin, Germany) was used. “Photon counting was performed with a picosecond resolution. GFP was excited with a 485 nm (LDH-D-C-485, 32 MHz, PicoQuant, Berlin, Germany) pulsed polarized laser. The laser power at the objective lens was adjusted to 1 µW. mCherry was excited with a 561 nm laser at 1% power. Light, emitted from the sample, was separated by a polarizing beam splitter and a LP610 beam splitter before photons were selected with a fluorophore-specific band-pass filter. Photons were detected in both donor and acceptor channels simultaneously with Tau-SPADs (PicoQuant, Berlin, Germany). Images were acquired at zoom 8 resolution of 256x256pixel with a pixel size of 0.1 µm and a pixel dwell time of 12.54 µs and laser repetition rate of 32 MHz. Photons were collected over 40 frames.

Before image acquisition, the system was calibrated. First, the objective was adjusted to reach a maximal count rate. Fluorescence-Correlation-Spectroscopy (FCS) curves of Rhodamine110 dye and water were acquired to monitor the system function. Internal Response Function (IRF) for each laser were determined by measuring the fluorescence decay of quenched erythrosine in saturated potassium iodide using the same hardware settings as for the FRET pair.

In order to determine the average lifetime of CRK18-GFP recombinant protein, regions of interest (ROIs) of the plasma membrane in the FLIM image were selected manually. The fluorescence decay times of the ROIs were analysed using SymPhoTime FLIM analysis software (SymPhoTime 64, version 2.4; PicoQuant, Berlin, Germany). Time-correlated single photon counting (TCSPC) bins of donor channel 1 (parallel light) and channel2 (perpendicular light) were binned 16 times. Pixels above the pile-up limit (10% of the lase repetition rate) and chloroplasts were removed manually. The data was fitted using the grouped FLIM image analysis tool (n-exponential reconvolution, n=2). The intensity-weighted lifetime was considered as the sample’s apparent lifetime. All samples were measured in at least two independent experiments.

### CRK18 dimer modelling and coarse-grained (CG) molecular dynamics simulations

The initial structural model of the CRK18 homodimer was generated using AlphaFold3 (Abramson et al. 2024). Structure prediction was performed using the ectodomain sequence of CRK18, excluding the signal peptide, corresponding to residues 25-249 of the CRK18 sequence (UniProt Q8RX80). The highest-confidence model, selected based on the predicted interface and overall structural confidence metrics (ipTM and pTM), was subsequently converted into a CG representation using the MARTINI 3 force field (Souza et al. 2021). To preserve the native tertiary and quaternary structure throughout the simulations, an elastic network was generated with Martinize2 (Kroon et al. 2025) using lower and upper distance cutoffs of 0 and 0.85 nm, respectively. The CG model was embedded in a cubic simulation box using a 14 Å padding distance with INSANE (Wassenaar et al. 2015), solvated with standard MARTINI water beads, and neutralized in the presence of 0.15 M NaCl.

All molecular dynamics simulations were performed with GROMACS 2024.5 (Abraham et al.) under periodic boundary conditions. The systems were first energy-minimized using the steepest descent algorithm until the maximum force was below 10 kJ mol⁻¹ nm⁻¹. An initial 2 ns equilibration was performed under NVT conditions using a 10 fs integration time step to stabilize the system temperature. This was followed by a 2 ns NPT equilibration employing the Berendsen barostat to allow gradual density and pressure relaxation while maintaining the 10 fs time step. Finally, a 10 ns NPT equilibration was performed using the same pressure-coupling scheme with a 20 fs integration time step, matching the production simulation conditions. Production simulations were subsequently carried out for 10 µs using a 20 fs integration time step under NPT conditions. Temperature was maintained at 298 K using the velocity-rescale thermostat, whereas pressure was controlled semi-isotropically at 1 bar with the Parrinello-Rahman barostat. Three independent simulations were initiated from the same initial structure using different randomly generated initial velocity seeds to improve conformational sampling.

### Backmapping and all-atom (AA) molecular dynamics simulations

Representative CG conformations were converted to atomistic resolution using the CG2AT backmapping (Vickery and Stansfeld 2021) with MARTINI 3 fragments and the CHARMM36 force field (Best et al. 2012). The resulting atomistic models were subsequently prepared using CHARMM-GUI (Park et al. 2023). Protonation states were assigned assuming an apoplastic pH of 5.0, and each system was solvated in a rectangular box with TIP3P water molecules using a 14 Å padding distance. Sodium and chloride ions were added to achieve electrical neutrality and a final NaCl concentration of 0.15 M. All-atom molecular dynamics simulations were performed with NAMD 3.0.2(Phillips et al. 2005) using the CHARMM36 force field (Best et al. 2012) under periodic boundary conditions. Long-range electrostatic interactions were treated using the particle-mesh Ewald (PME) method, whereas short-range van der Waals interactions were smoothly switched between 8 and 10 Å with a 12 Å pair-list cutoff. Each system was first energy-minimized by 5000 conjugate-gradient steps, followed by a 100 ps equilibration under NPT conditions at 298 K and 1 atm. Production simulations were subsequently performed for 200 ns under NPT conditions at 298 K and 1 atm. Temperature was controlled using Langevin dynamics, pressure was maintained with the Langevin piston method, and covalent bonds involving hydrogen atoms were constrained using the SHAKE algorithm.

### Trajectory and structural analyses

Trajectory analyses were performed using GROMACS 2024.5 (Abramson et al. 2024) and the MDAnalysis Python library v2.10.0 (Gowers et al., 2016). Prior to analysis, trajectories were structurally aligned to the initial simulation frame. CG protein structural stability was evaluated by calculating the root mean square deviation (RMSD) and RMSD distributions were used to identify the predominant conformational state for subsequent backmapping and structural analyses. For the AA simulations, the distance between the centers of mass of both monomers (COM-COM) were measured. RMSD distributions were used to identify the predominant conformational states sampled during the simulations, from which representative structures were extracted. Representative structures were further analyzed using PDBePISA (Krissinel et al., 2007) to characterize the dimer interface. The binding affinity of the representative dimer conformations and the crystallographic structure was estimated using PRODIGY(Xue et al. 2016) and PRODIGY-CRYSTAL(Elez et al. 2018). The pKa values of all cysteine residues were predicted using PROPKA3 (Olsson et al. 2011). To assess the contribution of dimerization to the predicted pKa values, chain A was extracted from the representative closed homodimer and analyzed separately as a monomeric structure. The electrostatic surface potential was calculated by numerically solving the non-linearized Poisson– Boltzmann equation using the APBS plugin implemented in Chimera 1.19 (Pettersen et al. 2004), with an ionic strength of 0.15 M NaCl, temperature of 298 K, and default dielectric parameters. Trajectory visualization was performed using VMD 2.0.0a7(Humphrey et al. 1996), whereas figures were generated with PyMOL (Schrödinger, LLC. The PyMOL Molecular Graphics System, Version 3.1.6.1).

### Generation of Arabidopsis transgenic lines and growth conditions

To obtain stable expression lines of CRK18 recombinant protein in Arabidopsis, genomic CRK18 DNA was amplified from Arabidopsis and cloned into the pGIIB-p35S-LIC_HpaIv2-3xFLAG-mScarlet-I-tNOS vector. The construct was transformed into *Agrobacterium tumefaciens*, and stable Arabidopsis lines in Col-0 ecotype in roGFP2-Orp1 background were generated using the floral dip method. Stable roGFP2-Orp1 expressing Col-0 was used as a control for the CRK18 overexpression lines.Expression and localisation of CRK18-3xFLAG-mScarlet-I in Arabidopsis leaves was checked using a Leica SP5 confocal microscope (Ex 561 nm, Em 595-645 nm).

Arabidopsis plants were grown under long day conditions (16h light - 8h dark, 21 °C, 60% humidity). Four-week-old rosettes were harvested, the tissue was ground to a fine powder and stored at -80 °C until use in IP-MS and (phospho)proteomics experiments.

### Total and Phosphoproteome measurements

To assess whether overexpression of CRK18 affects protein expression and signalling in Arabidopsis, we measured the total proteome and the phosphoproteome. For the measurements, four-week-old Arabidopsis rosettes were used of CRK18-OE1, CRK18-OE2, and the roGFP2-Orp1 background line as a control. Four replicates of each line were prepared and measured.

First, protein was extracted from the ground tissue samples. The tissue was resuspended in extraction buffer (100 mM Tris-HCl, pH 8.5, 7M Urea, 1% NP-40, 10 mM DTT, 10 U/mL DNase-I (Roche), MgCl2, 1% Benzonase (Novagen) and sonicated (Qsonica) for 10 minutes. Particles were pelleted by centrifugation for 20 min and discarded. Next, the protein extract was incubated with 1% benzonase and 20 mM iodoacetamide at 25 C for 30 min, followed by methanol chloroform precipitation of the protein. Triethyl ammonium bicarbonate was added, and the protein pellet was resuspended by sonication for 15 min. A BCA assay was performed to determine the protein concentration (Pierce). For phosphoproteomics experiments, 500 µg protein was used per sample, and 100 µg was used for total proteome samples. Trypsin was added to the samples (1:100 trypsin:protein ratio) and incubated ON at 25 C. 10% TFA was added to adjust the pH to 2-3 and the peptides were purified using C18 microcolumns. Columns were prepared by using C18 octadecyl discs (Empore) LiChroprep RP-18 (Merck), and equilibrated in 0.1% formic acid. The peptides were applied to the columns. The columns were washed with 0.1% formic acid, and peptides were eluted in 80% acetonitrile, 0.1% formic acid. Total proteome samples were not processed further, the acetonitrile was evaporated using a vacuum centrifuge. Samples were resuspended in 0.1% formic acid.

The phosphorylated peptides in the phosphoproteomics samples were enriched for with Fe-NTA MagBeads (Cube Biotech). Samples were diluted 1:1 in loading buffer (80% acetonitrile, 5% TFA, 0.2% Glycolic Acid), added to 100 µL equilibrated Fe beads, and incubated for 20 min. Beads were washed one time in loading buffer, one time in wash buffer 1 (80% AcNi, 1% TFA) and one time in wash buffer 2 (80% AcNi, 0.2% TFA). Phosphopeptides were eluted from the beads by incubating for 15 min in 100 µL 1% NH4OH. Next 10% formic acid was added to the elution to acidify the sample, and another C18 microcolumn step was performed. Phosphoproteome samples were also dried by vacuum centrifuge and resuspended in 0.1% formic acid.

Samples were measured using nano liquid chromatography–tandem mass spectrometry (nLC–MS/MS), and the data was analysed using MaxQuant and further processed in R as described previously (Kuhn et al. 2024). The mass spectrometry proteomics data generated during this study have been deposited in the MassIVE repository under accession number MSV000102194. The dataset is currently available to reviewers using the provided reviewer credentials and will be made publicly available upon publication.

### Immunoprecipitation – mass spectrometry

To detect possible interaction partners of CRK18, IP-MS experiments were performed. Two CRK18 Arabidopsis overexpression lines were used (35S:CRK18-3xFLAG-mScI #1 (CRK18-OE1) and 35S:CRK18-3xFLAG-mScI #2 (CRK18-OE2)), and a control of the roGFP2-Orp1 background. For each of the three lines, four replicates were measured. First protein was extracted from ground tissue by incubating for 30 minutes at 4 °C in extraction buffer (50 mM Tris-HCl pH7.5, 150 mM NaCl, 10% glycerol, 10 mM EDTA, 2 mM DTT, cOmplete EDTA-free protease inhibitor (Sigma), 1 % v/v IGEPAL). After incubation, particles were filtered from the protein extract, and anti-FLAG agarose beads were added (Chromotek). Protein extract was incubated with the beads for 2 hours at 4 °C. After incubation, the supernatant was discarded, and the beads were washed two times with wash buffer (20 mM Tris-HCl pH7.5, 150 mM NaCl), and two times with ammonium bicarbonate buffer. Next, the sample was reduced on the beads by incubating for 30 minutes at 45 C with 20 mM DTT in ammonium bicarbonate buffer. Reduction was followed by a 30-minute incubation with 20 mM iodoacetamide at room temperature, after which 20 mM cysteine was added. Samples were digested by overnight incubation with Trypsin. Beads were removed by centrifugation, and Trypsin was inactivated by the addition of 10 % TFA. Samples were measured by nano liquid chromatography-tandem mass spectrometry (nLC–MS/MS) and analysed by MaxQuant as described previously (Tyanova et al. 2016a; Feng et al. 2022). Filtering and statistical analysis of the data were performed with Perseus (Perseus V 1.6.2.1) (Tyanova et al. 2016b). Gene ontology (GO) term enrichment analysis was performed on the significantly enriched proteins identified in CRK18-OE1 and -OE2 relative to the list of identified proteins using the PANTHER GO web tool (Ashburner et al. 2000; Thomas et al. 2022; Consortium et al. 2023). The mass spectrometry proteomics data have been deposited to the ProteomeXchange Consortium via the PRIDE partner repository with the dataset identifier PXD080730 (Perez-Riverol et al. 2025).

### Single particle photoactivated localization microscopy (sptPALM) and photochromic reversion

VA-TIRFM imaging and photochromic reversion-enabled long-term sptPALM imaging were performed as previously described in (Arx et al. 2024). Briefly, VA-TIRFM was operated on a custom-build TIRFM set-up (Arx et al. 2024; Rohr et al. 2024)equipped with an 100x objective NA 1.49 (Zeiss, 421190-9800-000), four laser lines (405, 488, 561 and 642 nm), a polychromatic modulator (AOTF, AA OPTO-ELECTRONIC) and a sCMOS camera (Hamamatsu Photonics, ORCA-Flash4.0 V2). To ensure that the target laser power value always corresponds to the irradiance in the sample plane, the power of each laser line was calibrated before each experiment using a PM100D detector (Thorlabs). 3-week-old *N. benthamiana* leave samples (28 to 34 hours post Agrobacterium infiltration) were delicately mounted between two coverslips (Epredia 24×50 mm #1) in liquid ½ MS medium and placed on the specimen stage without additional weight. Images were acquired at 20 Hz frame rate (50 ms). For photochromic reversion, mEOS3.2 was photoconverted, excited and kept in a prolonged fluorescent state using 1000 µW 561nm laser power, 2-15 µW 405 nm laser power and 1-30 µW 488 nm laser power. The emitted light was collected using 568 LP Edge Basic Long-pass Filter, 584/40 ET Band-pass filters (AHF analysentechnik AG) and recorded from a 51.2 x 51.2 µm region of interest with 100 nm pixel size. Track reconstruction and analysis was carried out as previously described (Arx et al. 2024). To ensure bona-fide single-molecule tracking we analysed frames with relatively low molecule density (ca. 0.1 – 1 molecule per µm^2^). We used TrackMate7 (Ershov et al. 2022)in Fiji (Schindelin et al. 2012)to reconstruct single-particle trajectories. Single particles were segmented frame-by-frame by applying a Laplacian of Gaussian (LoG) filter and estimated particle size of 0.3 µm with a Quality threshold of 30. Individual single particles were localized with sub-pixel resolution using a built-in quadratic fitting scheme. Single-particle trajectories were reconstructed using a simple linear assignment problem (Jaqaman et al. 2008)with a maximal linking distance of 0.4 µm and a 2 frames-gap-closing maximum. The coordinates of the single particle trajectories were then further analysed using a custom-made python script to calculate the mean square displacement (MSD) and diffusion coefficient (D) in batch mode. Only tracks with a minimum of five localizations were kept for analysis. The MSD and diffusion coefficient was calculated based on the first four time points of each trajectory as previously described(Saxton and Jacobson 1997). To analyze spatial arrests, which occur within one track we used the previously published hybrid machine learning framework CASTA (Arx et al. 2026b)(https://pypi.org/project/casta/). Briefly, CASTA consists of a trained machine learning core classifier, which together with four diffusional signature metrics predict for timepoint to be either arrested or non-arrested based on a majority voting scheme. As an input, the folder containing the trackmate files containing coordinates (x, y) and timepoints (t) as well as the unique track ID were provided. From this the step length, MSD and diffusion coefficient (D) are calculated for each timepoint in a sliding window of ten timepoints, standardized and passed on to a trained Hidden Markov Model (HMM). Next, four diffusional features including the step length, successive angles, 2-dimensional kernel density estimation (KDE) as well as consecutive self-intersections are computed for each timepoint and compared to their respective threshold value, which was found by maximising the separation between the true positive and true negative class using simulated ground truth data. As an output CASTA automatically computes diffusional summary metrics such as the number of arrested and non-arrested timepoints, the diffusion of arrested and non-arrested molecules as well as the duration and area of arrestment. According to the imaging conditions, the frame rate was set to 50 ms and the minimum required track length was set to the default value of 25 timepoints.

### Nematode sterilization and in vitro inoculation

*Heterodera schachtii* (Woensdrecht population from IRS, The Netherlands) cysts were obtained from infected *Brassica oleracea* roots grown in sand (Baum et al. 2000). The cysts were hatched for 7 days in a solution containing 1.5 mg · ml^-1^ gentamicin sulfate, 0.05 mg · ml^-1^ nystatin, and 3 mM ZnCl2. Next, *H. schachtii* second-stage juveniles (J2s) were separated from debris using a 35% sucrose gradient and incubated in a sterilization solution (0.16 mM HgCl2, 0.49 mM NaN3, and 0.002% Triton X-100) for 15 minutes. Finally, the J2s were washed three times with sterile tap water and re-suspended in 0.7% Gelrite (Duchefa Biochemie, Haarlem, The Netherlands). For the in vitro infection assays, 10 µl of a gelrite solution containing ≈250 J2s was inoculated onto the roots of 14-day-old Arabidopsis plants grown in Knop medium (Sijmons et al. 1991)at 21°C under 16 h: 8 h, light: dark conditions, in 12-well cell culture plates (Greiner bio-one). The number of J3 males and/or cysts was counted at 14 and 28 dpi.

## Results

### Cysteines are important for the stability of the CRK18-ECD *in vitro*

To investigate the effect of cysteines on the stability of CRK18 ECD, we selected three of the six predicted disulphide bonds in the CRK18-ECD and generated cysteine-to-alanine mutants in which both cysteines were substituted (Figure 1A). C94-C122 and C216-C241 were selected to represent disulphide bonds within the conserved motifs of DUF26-A and DUF26-B, respectively. C227-C228 was chosen as a variable, clade-specific predicted disulphide bond formed by two vicinal cysteines, which is rare in proteins (Richardson et al. 2017a; Stouthamer et al. 2026). We purified the ECD of CRK18 and its mutant variants using the baculovirus system in TniH5 insect cells. All variants behaved similarly during purification and were obtained with high purity (Supp. Figure 1A-D). We used SEC-MALS conjugate analysis to determine the molar mass, oligomerisation and glycosylation state of CRK18-ECD^WT^ and a cysteine mutant variant, CRK18-ECD^C227A,C228A^. In SEC-MALS experiments, the chromatograms of both protein variants showed a single main peak with a small, unquantifiable peak to its left (Supp. Figure 1E). We characterised the proteins from the main peak as a monomer with a total mass of 34 kDa, of which 6 kDa corresponds to glycosylation and the other 28 kDa to the protein (Supp. Figure 1E). The experimentally determined MW of the protein without glycosylation (28 kDa) is in line with the predicted MW of 29.3 kDa based on the amino acid sequence.

**Figure 1.**
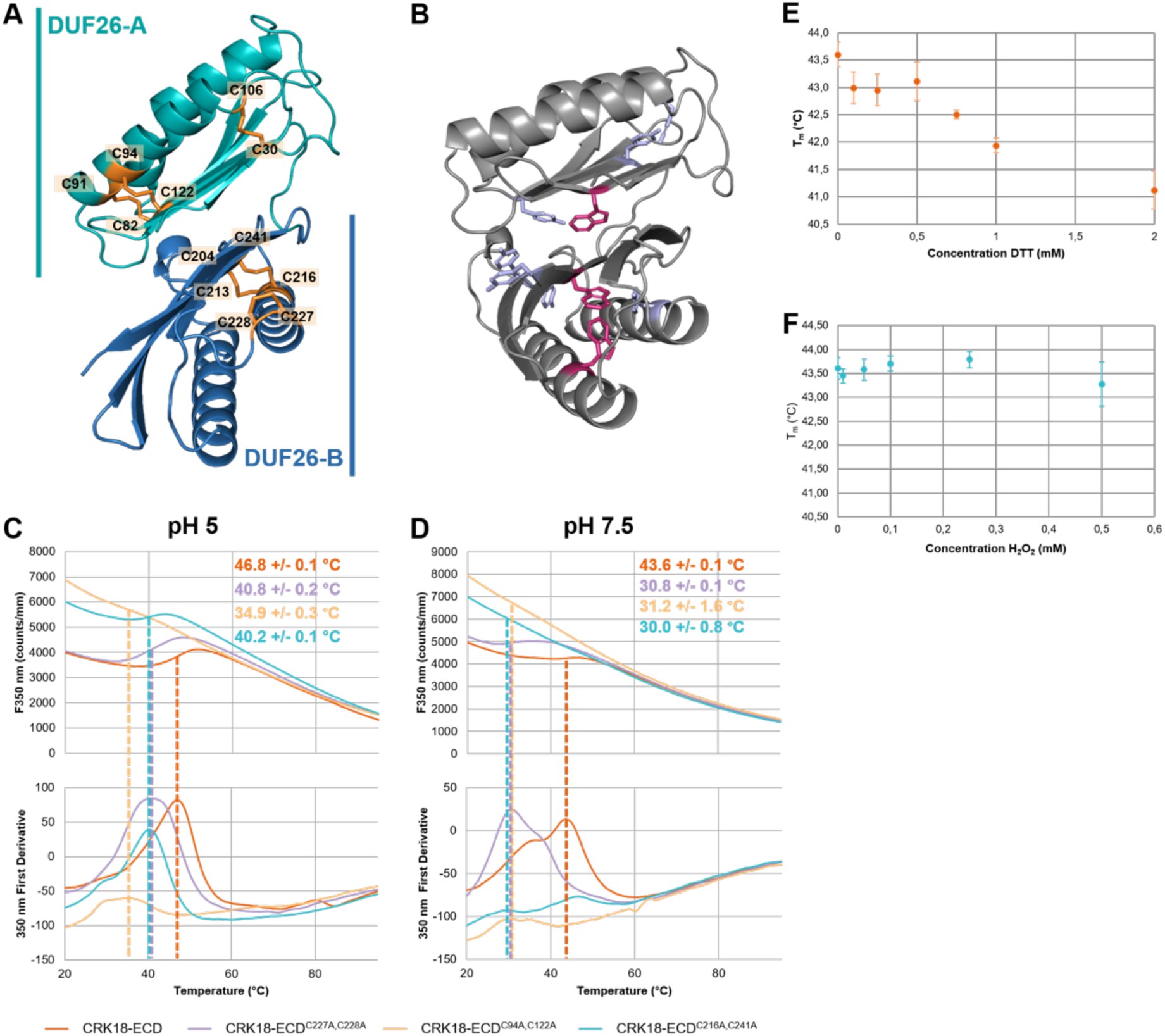
CRK18-ECD cysteine mutant variants have decreased thermal stability compared to CRK18-ECD. A) AlphaFold model of CRK18-ECD. A cartoon representation of the ECD is shown with the DUF26-A and DUF26-B domains marked in cyan and blue, respectively, and the cysteines coloured in orange. B) AlphaFold model of CRK18-ECD with tryptophans and tyrosines coloured in magenta and light blue, respectively. The thermal unfolding of CRK18-ECD and mutant variants was measured at C) pH 5 and D) pH 7.5 using nanoDSF. Graphs show thermal unfolding curves of fluorescence emission at 350 nm as a function of temperature (top) and its first derivative (bottom). The thermal midpoint of unfolding, Tm, indicated by the dashed vertical lines, was determined by deriving the inflection point (IP) of the first derivative of the thermal unfolding curve. Tm-values and corresponding standard deviations (SD) are marked in the top right. Note that at pH 7.5 the first derivative of the fluorescence emission at 350 and 330 nm also shows a peak at approximately 36°C besides the peak at 43.6 °C, indicating two subsequent unfolding events. All samples were measured in triplicate in standard capillaries. pH 5 samples were prepared in 20 mM NaOAc, 150 mM NaCl, and 0.1% Pluronic acid. pH 7.5 samples were prepared in 20 mM HEPES, 150 mM NaCl, 0.1% Pluronic acid. E-F) DTT and H_2_O_2_ impact on the thermal stability of CRK18-ECD. nanoDSF measurements were performed on CRK18-ECD at different DTT and H_2_O_2_ concentrations. All samples were prepared at a protein concentration of 0.4 mg/ml in 20 mM HEPES, 150 mM NaCl, 0.1% Pluronic acid pH7.5, at varying concentrations of DTT (0-2mM) and H_2_O_2_ (0-0.5 mM). From the temperature-dependent 350 and 330 nm fluorescence intensities and the ratio thereof the temperature at which 50% of the protein is unfolded (Tm) was determined. Graphs present Tms of CRK18-ECD at different H_2_O_2_ (E) and DTT (F) concentrations.

We assessed the thermal stability of the CRK18-ECD using nano differential scanning fluorometry (nanoDSF), which measures changes in intrinsic fluorescence of tryptophan (Trp) and, to a lesser extent, tyrosine (Tyr) upon temperature-induced protein unfolding(Martin-Ramirez et al. 2026). From this change in fluorescence, the melting temperature (T_m_), or thermal midpoint, at which 50% of the protein molecules are unfolded, can be determined. The CRK18-ECD contains three Trp residues, of which two are buried in DUF26-B, and one resides in the interface between both DUF26 domains, as well as seven Tyr residues that are largely buried (Figure 1B). The T_m_ values of CRK18-ECD were determined at pH 5 and 7.5 to cover the dynamic pH range of the apoplast. Thermal unfolding curves recorded at 350 nm show that the T_m_ of CRK18-ECD at pH 5 is 46.8 ± 0.1°C, and at pH 7.5, it is 43.6 ± 0.1°C (Figure 1 C and D, Supp. Figure 2). The increased protein stability of the CRK18-ECD at pH 5 compared to pH 7.5 may reflect its biological environment in the cell, as the pH of the apoplast is reported to be approximately 5 under resting (basal) conditions^26^. An increase in temperature typically leads to protein unfolding, measured by a decrease in fluorescence intensity in nanoDSF due to increased exposure of Trp to the solvent. Interestingly, the thermal unfolding curves of CRK18-ECDs showed an increase in fluorescence between ∼40 and 50 °C, suggesting that Trp fluorescence is quenched in native CRK18-ECD, possibly due to the presence of disulphides (Hennecke et al. 1997).

**Figure 2.**
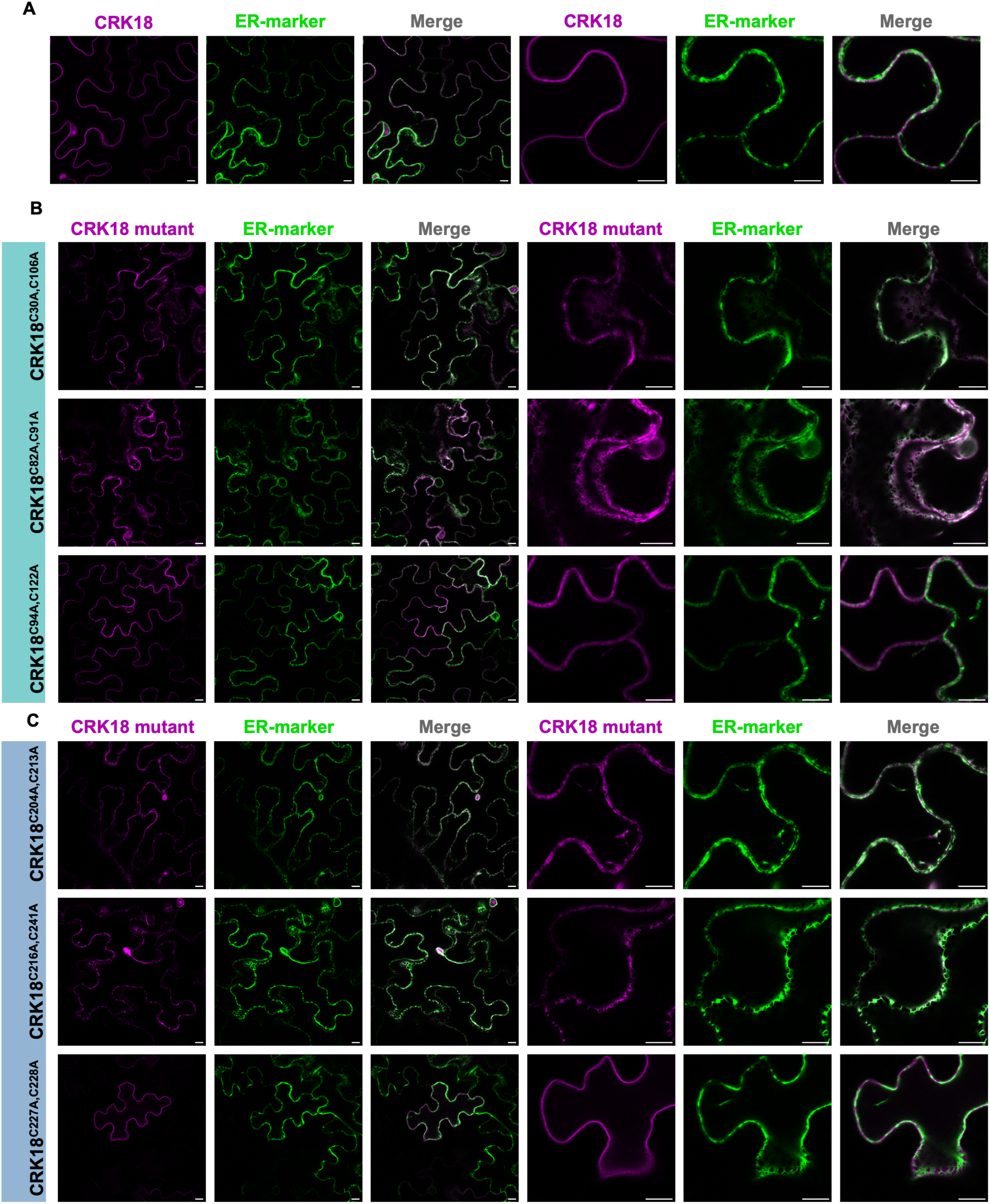
Most of the cysteine residues are required for the folding and localization of CRK18 to the plasma membrane. A) Transient expression of 35S:CRK18^K367N^-3xFLAG-mScI (CRK18) and 35S:AtVMA21a-mNG (ER-marker) in N. benthamiana leaf epidermal cells. Images show an optical section through the cell center. B) Co-Localization of cysteine mutants in DUF26-A; CRK18^C30A,C106A^, CRK18^C82A,C91A^ and CRK18^C94A,C122^ with the ER marker. C) Co-localization of DUF26-B cysteine mutants CRK18^C204A,C213A^, CRK18^C216A,C241A^ and CRK18^C227A,C228A^ with the ER marker. CRK18^C227A,C228A^ does not co-localize with the ER-marker. CRK18 and all Cysteine-to-Alanine mutant variants included the K367N mutation and were expressed under a 35S promoter with C-terminal 3xFLAG-mScarlet-I tags. Scale bars are 10 µM.

CRK18-ECD cysteine mutants have lower thermal stability than CRK18-ECD (Figure 1C and D). At pH 5, both CRK18^C227A,C228A^-ECD and CRK18^C216A,C241A^-ECD have T_m_-values ∼40 °C, which is a decrease in T_m_ compared to CRK18-ECD. The stability of these mutant variants also depends on pH, with the mutants being more stable at pH 5 than at pH 7.5 (Figure 1C and D, Supp. Figure 2). However, the increase in stability upon lowering pH is much higher for the mutant variants than for CRK18-ECD. The T_m_ of CRK18-ECD differs by ∼3 °C between pH 7.5 and 5, whereas for CRK18^C227A,C228A^-ECD and CRK18^C216A,C241A^-ECD, the Tm difference is ∼10 °C. This pH-dependency is not apparent in CRK18^C94A,C122A^-ECD. The unfolding curves of CRK18^C94A,C122A^-ECD at both pH values show a broad unfolding transition, and the corresponding T_m_s are difficult to determine and are estimated at ∼30 °C, indicating that CRK18^C94A,C122A^-ECD is rather unstable.

The specificity of the CRK18-ECD interactions was shown to depend on the presence of ROS (Martin-Ramirez et al. 2026). To test whether CRK18-ECD stability is affected by changing redox status, we measured its T_m_ after treatment with H_2_O_2_ or DTT (Figure 1E and F, Supp. Figure 3, 4). Treatment with different concentrations of H_2_O_2_ did not affect the thermal stability of CRK18-ECD (Figure 1E, Supp. Figure 3). The cysteines could already be involved in disulphide bonds or are otherwise unaffected by the H_2_O_2_ treatment. Treatment with increasing DTT concentrations decreases the T_m_ of CRK18-ECD, which is likely caused by the reduction of disulphide bonds important for structural integrity (Figure 1F, Supplementary Figure 4).

**Figure 3.**
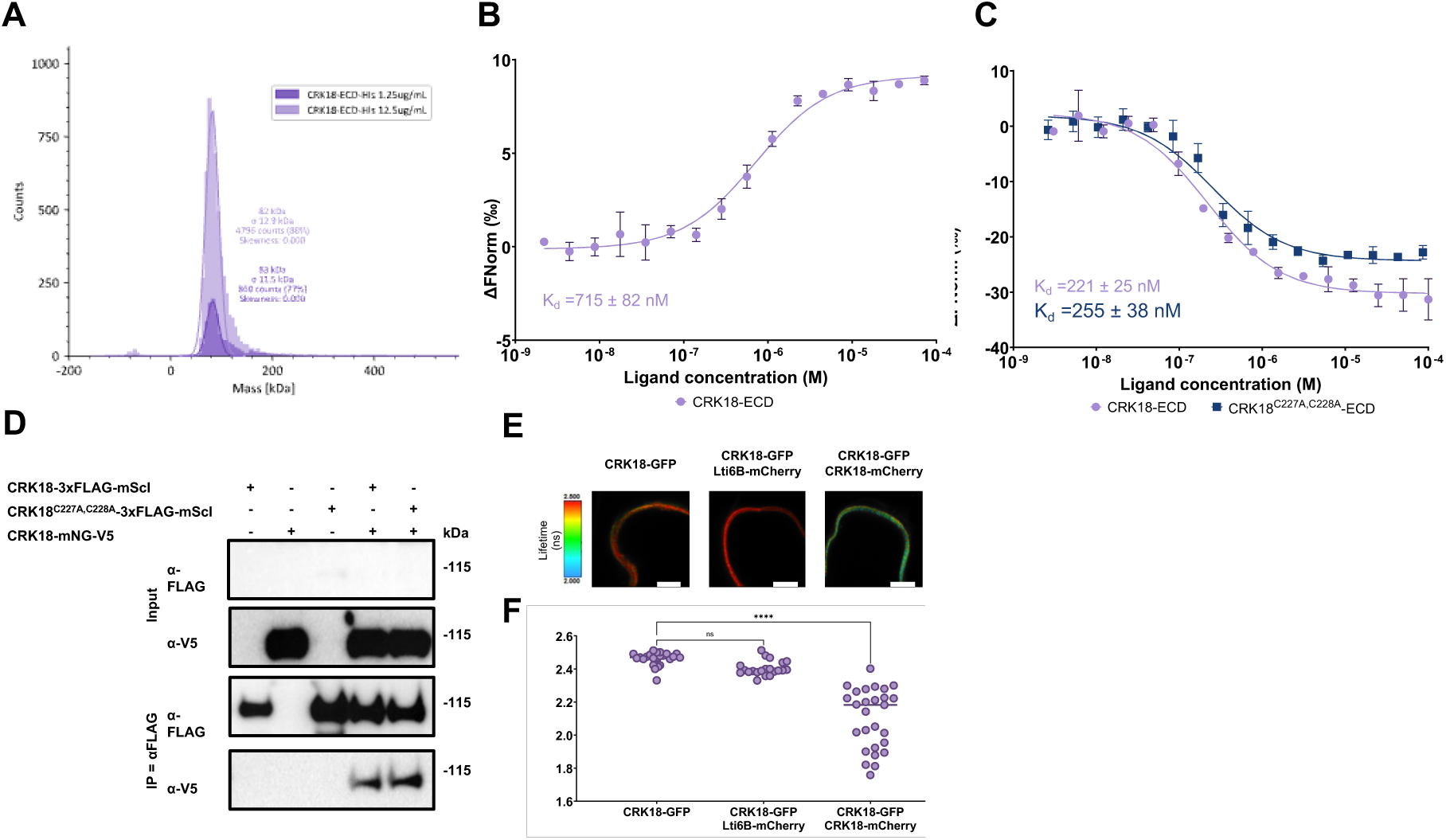
CRK18 homodimerizes on the level of the ECD and full-length receptor. A) Determination of particle sizes in S2 sample with Mass Photometry. Due to the ∼50 kDa detection limit of MP, monomeric CRK18-ECD could not be measured. Particles of ∼80 kDa, likely corresponding to dimers, were measured at two concentrations of CRK18-ECD. MP samples were diluted in NaOAc buffer pH5. B) Dose-response curve of labelled CRK18-ECD (target) with unlabelled CRK18-ECD (ligand) at pH 5. C) Dose-response curves of CRK18-ECD homodimerization and CRK18^C227A,C228A^-ECD homodimerization at pH 7.5 (n=3). For all MST measurements, a dilution series of unlabelled ligand was prepared, and labelled target was added at a constant concentration of 20 nM. The normalised change in fluorescence (ΔFNorm, ‰) was plotted against ligand concentration, from which the binding affinity (Kd ± SEM) could be determined. Samples at pH 7.5 were prepared in 20 mM HEPES, 150 mM NaCl, 0.1% Pluronic acid; samples at pH 5 were prepared in 20 mM NaOAc, 150 mM NaCl, 0.1% Pluronic acid. MST infrared laser power was set to 40% for all experiments. Excitation power varied: (B) 50%, (C) 100%. D) Co-immunoprecipitation of CRK18-mNG-V5 with CRK18^K367N^-3x-FLAG-mScI or CRK18^C227A,C228A,K367N^-3xFLAG-mScI visualised by western blot. *N. benthamiana* leaf tissue transiently expressing CRK18-FLAG and CRK18-V5 was harvested 2 days post-infiltration. Co-IP experiments were performed three times with similar results for CRK18. For CRK18^C227A,C228A^ one repeat showed less protein compared to CRK18. All 35S: CRK18 constructs contained the K367N mutation. The expected size of the recombinant protein without glycosylation is ∼103 kDa. E) CRK18 homodimerization was measured with FRET-FLIM. Representative confocal microscopy images of CRK18-GFP expressed at the plasma membrane in *N. benthamiana* cells. Images are pseudo-coloured based on the measured GFP average lifetime (ns) and represent cells expressing CRK18-GFP, CRK18-GFP-mCherry and co-expression of CRK18-GFP with Lti6B-mCherry (negative control) or CRK18-mCherry. Images were collected 8-12 hours after induction with estradiol. Scale bars are 6 µm. F) Quantified average lifetimes (ns) of GFP show a decrease upon co-expression of CRK18-GFP with CRK18-mCherry, indicating dimerization. The average lifetime was quantified for 21-27 repeat measurements. Statistically significant groups were determined by one-way ANOVA and Dunnett’s multiple comparisons test (ns= nonsignificant, **** = p<0.0001).

**Figure 4.**
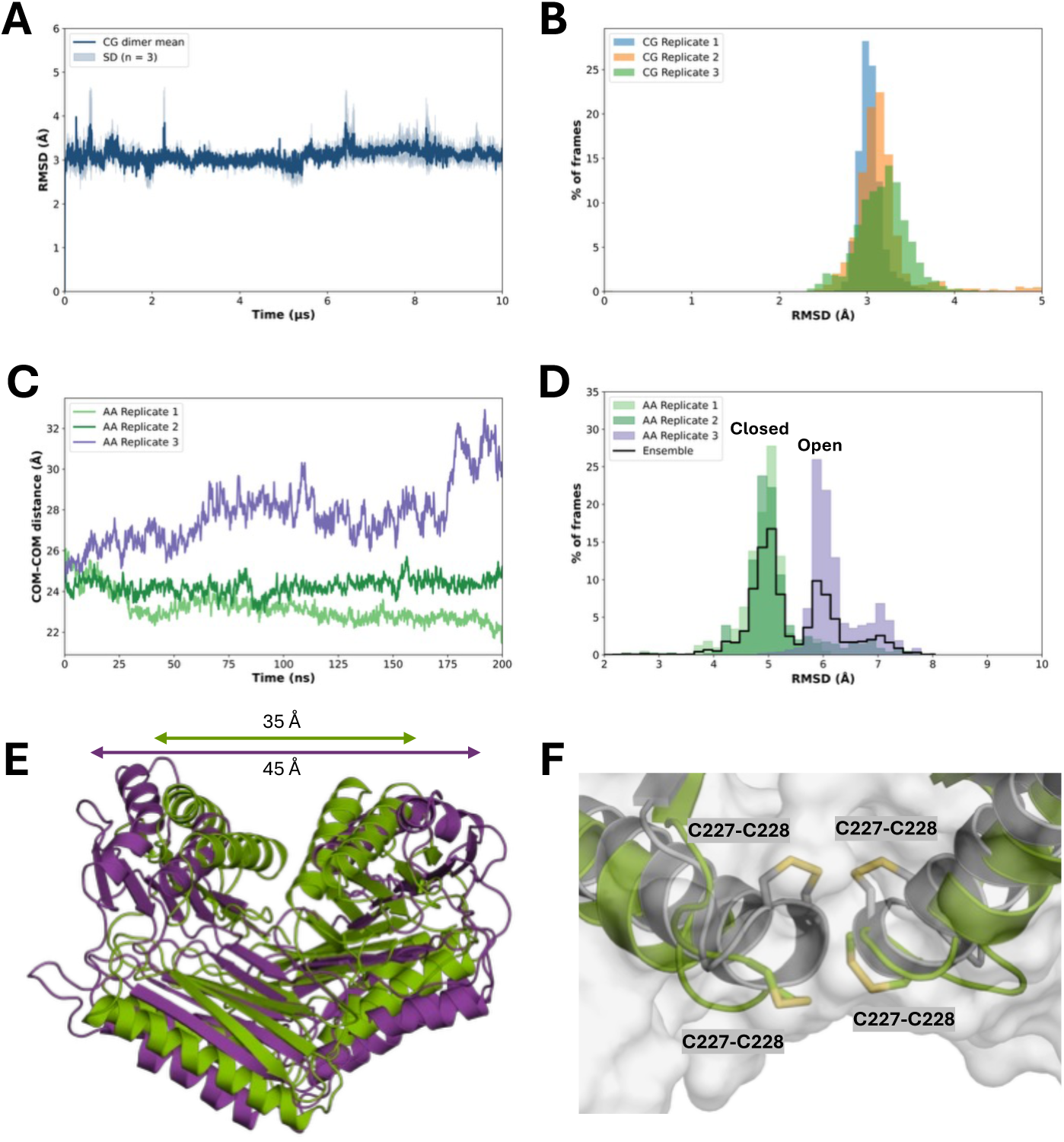
Multiscale molecular dynamics simulations identify a predominant closed conformation of the CRK18 ECD homodimer. A) Average backbone RMSD of the CG simulations over 10 µs, shown as the mean ± SD of the three independent replicas, indicating overall structural stability of the dimer. B) RMSD distribution of the CG trajectories, revealing convergence towards a predominant conformational state. C) Distance between the COM of the two monomers during the AA simulations. Replicates 1 and 2 remain in a compact arrangement, whereas replicate 3 samples a more extended configuration while the dimer remains associated. D) RMSD distributions of the AA trajectories, showing convergence of replicates 1 and 2 to the predominant closed conformation and sampling of a more open configuration in replicate 3. E) Superposition of representative closed (green) and open (purple) conformations extracted from the AA simulations, illustrating the increase in end-to-end distance from approximately 35 Å to 45 Å. F) Superposition of the initial AF3 model (grey) and the closed conformation from AA molecular dynamics simulations (green), showing the reorientation of the conserved C227–C228 pairs and their increased solvent exposure in the final state.

### Cysteines are important for localization of the CRK18 full-length protein

To assess how cysteines may affect CRK18 folding and function *in vivo*, we expressed the full-length proteins of CRK18 and a range of cysteine mutant variants in *N. benthamiana* and studied their cellular localization. Overexpression of CRKs has been shown to induce cell death in Arabidopsis and *N. benthamiana*(de Oliveira et al. 2016; Yadeta et al. 2017b). Transient expression of kinase-active full-length CRK18 under the constitutive 35S promoter, *35S:CRK18-3xFLAG-mScarlet-I* (mScI), in *N. benthamiana* leads to low protein expression (Supp. Figure 5) and cell death. As reported by Yadeta et al. (Yadeta et al. 2017b) for CRK28, mutating the conserved lysine (K367) in the kinase ATP-binding site to asparagine abolishes the cell death response in *N. benthamiana*, likely due to inactivation of the kinase domain. Targeting the corresponding lysine residue in CRK18 (CRK18^K367N^) increased expression levels (Supp. Figure 5). Thus, CRK18^K367N^ was used for all subsequent localization measurements of CRK18 and cysteine mutant variants in *N. benthamiana.* We observed that CRK18^K367N^-mScI marked the cell outline, consistent with its predicted plasma membrane (PM) localization. Furthermore, CRK18^K367N^-mScI localization followed a retracted PM pattern and marked Hechtian strands, cell wall-bound PM strands, upon plasmolysis (Supp. Figure 6A and B).

**Figure 5.**
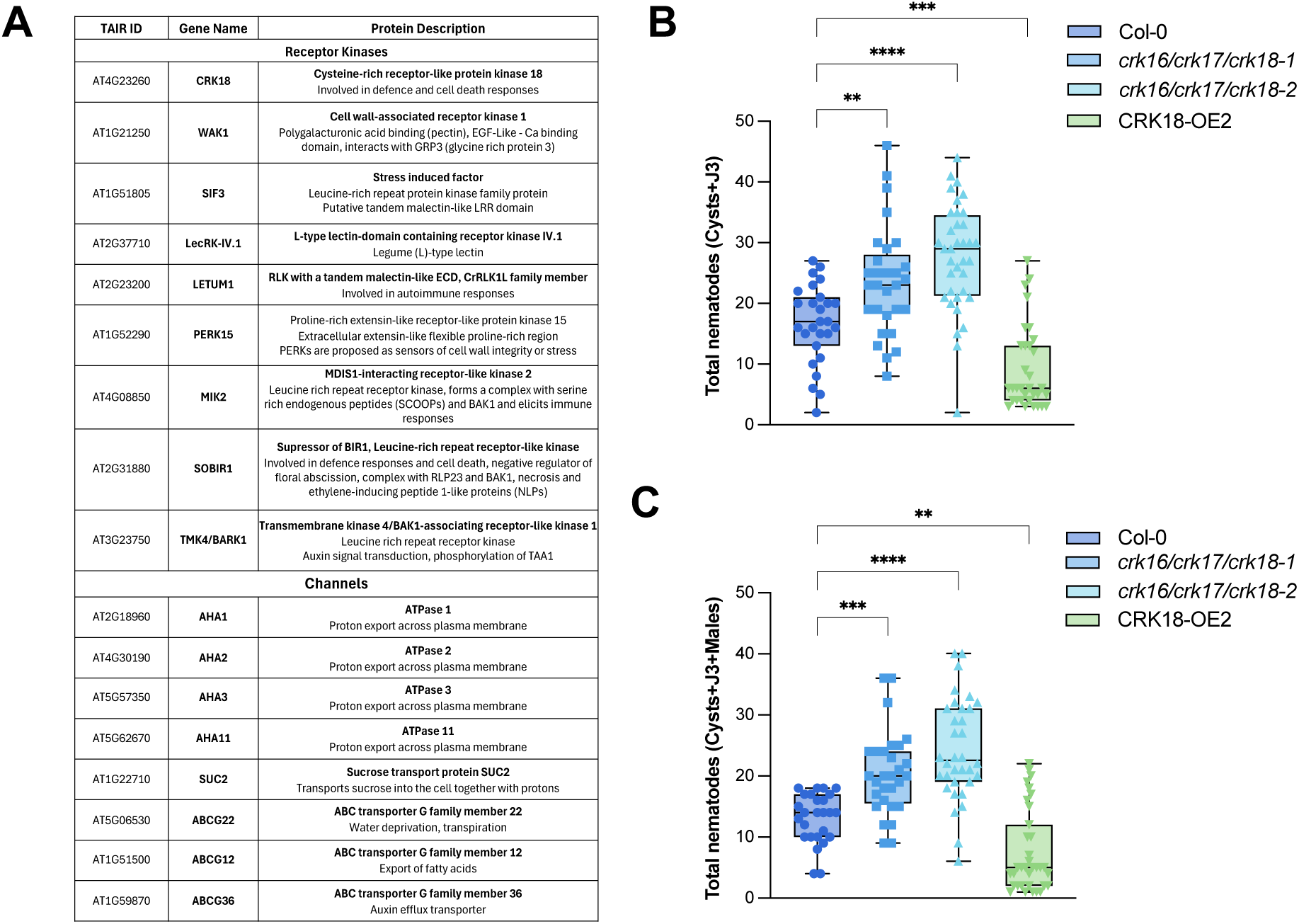
CRK18 plays a role in plants’ stress responses. A) CRK18 associates with other receptor kinases involved stress perception. List of significantly enriched (p-value < 0.01, log2(FC)> 1) plasma membrane-localized proteins identified by IP-MS of CRK18 in CRK18-OE1 and CRK18-OE2 in four-week-old Arabidopsis rosettes. CRK18 modulate plant susceptibility to the cyst nematode *Heterodera schachtii.* Two-week-old Arabidopsis seedlings were inoculated with ∼250 H. schachtii juveniles. B) The number of infections was counted 14 days post-inoculation. The graph shows the total number of cysts and J3 (developing juveniles). C) The number of infections was counted 28 days post-inoculation. The graph shows the total number of cysts, J3 (developing juveniles) and males. Each experiment was performed three times with similar results (n=25-36), which were pooled and analyzed using One-way ANOVA followed by Dunn’s post-hoc test with Benjamini-Hochberg correction for multiple comparisons (n=25-36).

**Figure 6.**
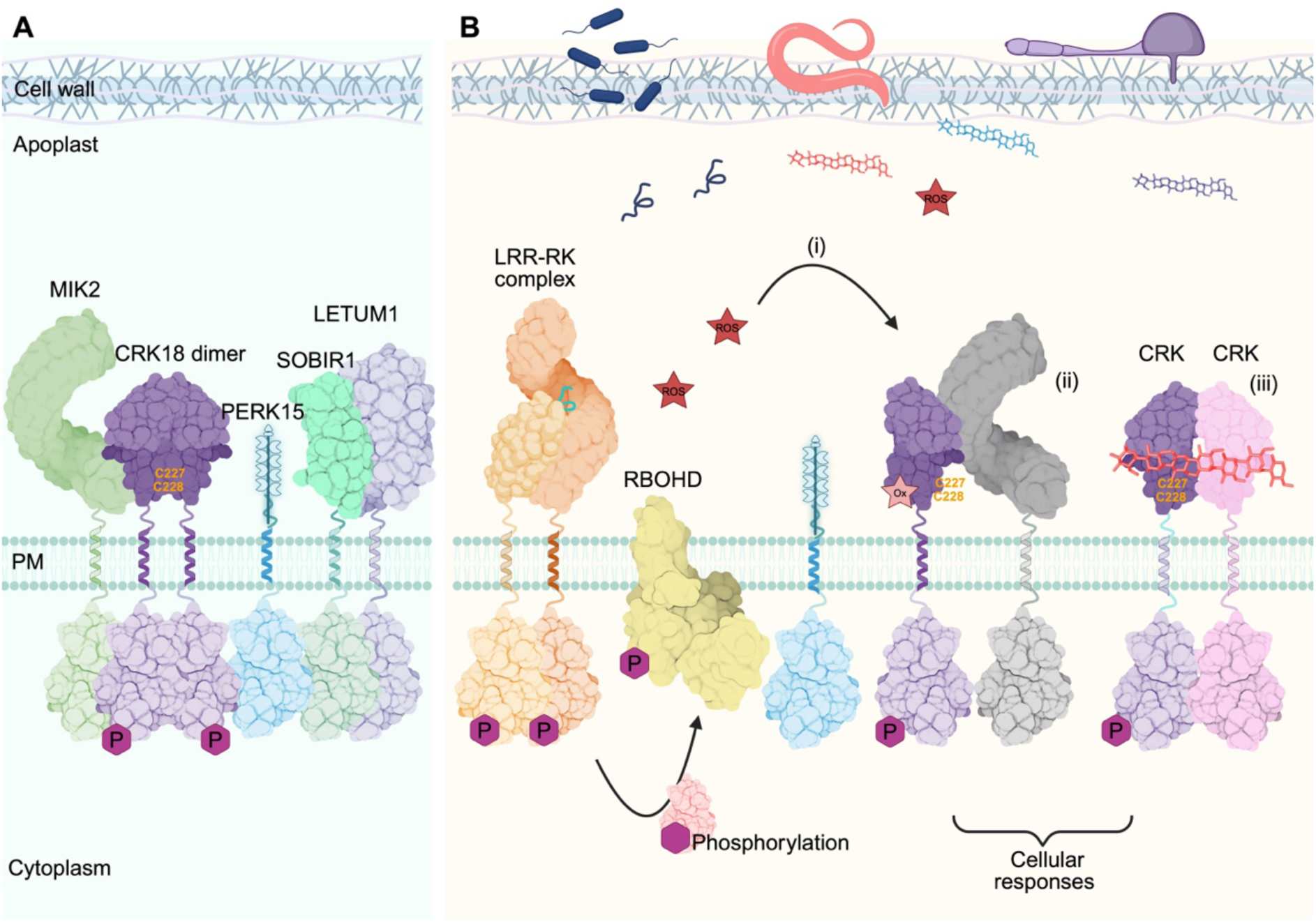
Potential roles of CRK18 in stress responses. A) In a basal state, CRK18 homodimers may associate with other receptor kinases (RKs) at the membrane. B) Upon perception of a stimulus, CRK18 activity and interaction partners could be modulated in several ways: i) CRK18, through their non-conserved cysteine residues, may be oxidatively modified by ROS produced by RBOHD, which is activated following phosphorylation events initiated by ligand-induced RK complex formation. ii) CRK18 may associate with other RKs, due to nanodomain reorganization. iii) CRKs may bind other ligands, such as glycans derived from the plant or microbial cell wall. Further research should be conducted to determine which stimuli activate CRKs and how this affects downstream signalling responses.

We designed cysteine mutant variants of CRK18 and tested their expression in *N. benthamiana*. The AlphaFold (AF) model of the CRK18-ECD predicts that all 12 cysteines are involved in disulphide bonds, including a bond between the sequentially adjacent cysteines C227 and C228 (Figure 1A) (Richardson et al. 2017a). For each of the 6 predicted disulphides in CRK18, double cysteine mutants were generated in which both cysteines were mutated to alanine (*35S:CRK18^C30A,C106A^-3xFLAG-mScI*, *35S:CRK18^C82A,C91A^-3xFLAG-mScI*, *35S:CRK18^C94A,C122A^- 3xFLAG-mScI*, *35S:CRK18^C204A,C213A^-3xFLAG-mScI*, *35S:CRK18^C216A,C241^-3xFLAG-mScI*, and *35S:CRK18^C227A,C228A^-3xFLAG-mScI*). While CRK18 localized uniformly at the PM, most CRK18 cysteine mutants exhibited an intracellular patchy localization pattern, reminiscent of the endoplasmic reticulum (ER) (Figure 2A). To test this, CRK18-mScI mutants and the ER-marker *35S:AtVMA21a-mNeonGreen* (mNG) were transiently co-expressed in *N. benthamiana*(Neubert et al. 2008). The ER forms an intracellular mesh-like network visible as a discontinued structure at the cell periphery in optical cross sections (Figure 2A). Several CRK18 cysteine mutants, including CRK18^C30A,C106A^, CRK18^C82A,C91A^, CRK18^C204A,C213A^ and CRK18^C216A,C241A^, lost strict PM localization and partially co-localized with the ER-marker (Figure 2B and C), suggesting that they are retained in the ER and to do not properly traffic to the PM. The localization pattern of CRK18^C94A,C122A^ is more uniform than that of the other cysteine mutants; suggesting that the protein is only partially retained in the ER (Figure 2B). Interestingly, CRK18^C227A,C228A^ has an even localization pattern at the PM, similar to wild-type CRK18 (Supp. Figure 6A and C). In addition, plasmolysis of CRK18^C227A,C228A^-mScI expressing tissue shows the presence of Hechtian strands, indicating that CRK18^C227A,C228A^-mScI localises to the PM (Figure 2C, Supp. Figure 6C and D). The CRK18^C227A,C228A^ mutant retains correct localization in the plasma membrane (Figure 2) and shows only a slight reduction in stability compared to the CRK18 (Figure 1). AlphaFold models suggest that C227 and C228 form a disulphide bond. Additionally, C227 has the highest solvent accessibility of all cysteines. This suggests that the vicinal disulphide between C227 and C228 may have a functional role, potentially as a redox switch (Supp. Table 1).

### CRK18 ECD homodimerizes *in vitro*

To elucidate further if cysteines contribute to CRK18 homodimer formation, we first confirmed CRK18 ECD dimerization previously presented in (Martin-Ramirez et al. 2026). To this end, we analysed the CRK18-ECD oligomeric state using mass photometry (MP), which infers particle size by measuring the amount of interference caused by the scattered light from particles landing on a glass slide and the internally reflected light (Foley et al. 2021). Indeed, particles of approximately 80 kDa were observable, corresponding to the approximate size of a CRK18-ECD dimer (Figure 3A).

Next, we assessed the binding kinetics of CRK18-ECD using microscale thermophoresis (MST). To measure CRK18-ECD homodimerization, we prepared samples with fluorescently labelled CRK18-ECD (target) and a dilution series of unlabelled CRK18-ECD (ligand). We measured CRK18-ECD homodimerization at pH 5 and 7.5 to approximate apoplastic pH under optimal growth and stress conditions. The interaction of CRK18-ECD to CRK18-ECD had K_d_-values of ∼700 ± 82 nM and ∼221 ± 25 nM at pH 5 and pH 7.5, respectively (Figure 3B and C). Overall, CRK18 homodimerization affinity remains in a similar range at both pH values, which suggests that pH-dependent changes in the apoplast may not strongly influence dimerization strength.

As mutating vicinal cysteines C227 and C228 did not substantially affect the stability of the CRK18-ECD and did not cause mislocalization of the full-length protein, we purified and performed MST on the CRK18^C227A,C228A^-ECD mutant. The vicinal cysteines are not conserved among CRKs and are likely candidates for the proposed role as ROS sensors. CRK18^C227A,C228A^-ECD interacted with CRK18^C227A,C228A^-ECD with a K_d_ of and ∼255 ± 38 nM at pH 7.5 (Figure 3C). The affinity of CRK18^C227A,C228A^ -ECD dimerization is similar to that of CRK18-ECD at pH 7.5 (Kd ∼221 nM), suggesting the vicinal cysteines do not directly (in the tested conditions) influence CRK18-ECD homodimerization.

Previously, we demonstrated that interfamily CRK dimerization is largely modulated by ROS (Martin-Ramirez et al. 2026). Under the conditions of the RIA assay, CRK18 dimerization was observed to be lower upon ROS treatment. To simulate changes in redox conditions *in vitro*, we treated CRK18-ECD MST samples with 100 µM H_2_O_2_ or 0.5 mM DTT at pH 7.5. H_2_O_2_-treated CRK18-ECD self-associates with a K_d_ in the high nm range (∼434 nM), similar to CRK18 without treatment (Supp. Figure 7). Under these conditions, H_2_O_2_ does not directly affect the interaction of CRK18-ECD with itself. The DTT-treated CRK18-ECD interacted with CRK18-ECD with a K_d_ of ∼580 ± 130 nM (Supp. Figure 7). Using the MST approach, neither H_2_O_2_ nor DTT treatment affected the homodimerization of CRK18-ECD, suggesting that the introduced changes in the redox environment do not have a major impact on the CRK18-ECD dimerization at pH 7.5.

### CRK18 homodimerizes *in vivo*

To assess CRK18 homodimerization and the influence of vicinal cysteines on this process at the level of the full-length receptor, we used co-immunoprecipitation (Co-IP) and Förster resonance energy transfer - fluorescence-lifetime imaging microscopy (FRET-FLIM).

To investigate the self-association of CRK18 with Co-IP, we transiently expressed *35S:CRK18^K367N^-3xFLAG-mScI* and *35S:CRK18^K367N^-mNG-V5* in *N. benthamiana*. We confirmed the localisation of the different recombinant versions of CRK18 to the plasma membrane by plasmolysis (Supp. Figure 8B). CRK18-3xFLAG-mScI was immunoprecipitated with anti-FLAG beads, and the presence of CRK18-mNG-V5 was detected via immunoblotting with anti-V5 antibodies, confirming CRK18 homodimerization (Figure 3D).

To examine the influence of the vicinal cysteines on homodimerization, we performed Co-IP of CRK18^C227A,C228A,K367N^-3xFLAG-mScI to CRK18^K367N^-mNG-V5 (Figure 3D). In one of three replicates, the mutant showed less protein in the pull-down compared to the wild type, but in the other two replicates, similar amounts of protein were detected (Figure 3D, Supp. Figure 8C). These results suggest the vicinal cysteines do not directly affect the homodimerization of CRK18. However, it is possible that the vicinal cysteines play a role in modulating the interaction upon perception of a specific signal.

Next, we used FRET-FLIM to measure CRK18 homodimerization. FRET-FLIM measures the decrease in the lifetime of a donor fluorophore (GFP) due to energy transfer to an acceptor (mCherry) when they are in close proximity to each other. Kinase-active CRK18-GFP and CRK18-mCherry were expressed transiently in *N. benthamiana* under an estradiol-inducible promoter. The fluorescence lifetime of CRK18-GFP was measured as 2.46 ± 0.04 ns in the absence of an acceptor and decreased in the presence of CRK18-mCherry to 2.10 ± 0.18 ns (Figure 3E, F). The negative control, Lti6B- mCherry, a small plasma membrane protein, did not decrease the GFP lifetime (2.40 ± 0.04 ns). The decrease in GFP lifetime of CRK18-GFP in the presence of CRK18-mCherry, and not in the presence of the negative control, indicates that CRK18 homodimerizes. Additionally, we measured FRET-FLIM using the kinase-inactive mutant, CRK18^K367N^, and observed a similar decrease in GFP lifetime (2.5 ± 0.02 ns to 2.2 ± 0.13 ns), indicating the kinase-inactive mutant does not affect dimerization of CRK18 (Supp. Figure 8A). Together, the results from Co-IP and FRET-FLIM experiments indicate that CRK18 homodimerizes on the level of the full-length receptor, and likely C227 and C228 do not directly modulate this interaction.

### Multiscale molecular dynamics simulations reveal a stable CRK18 ECD homodimer with a solvent-exposed vicinal cysteine pair

To obtain structural insight into CRK18-ECD homodimerization and the potential role of the C227-C228 vicinal cysteine pair, we performed multiscale molecular dynamics simulations starting from the highest-confidence AlphaFold3 (AF3) model of the CRK18 ECD homodimer (Abramson et al., 2024). The predicted dimer displayed a high-confidence interface (ipTM = 0.76) and quaternary structure (pTM = 0.84). Local confidence was high across most of the structure (pLDDT > 90), and the Predicted Aligned Error (PAE) map showed low alignment errors within and between monomers, supporting the reliability of the predicted assembly (Supp. Figure 9A, B).

The AF3 model was converted into a MARTINI coarse-grained (CG) representation (Supp. Figure 10A) and subjected to three independent 10 µs CG molecular dynamics simulations. In all replicates, the dimer remained associated and displayed only minor deviations from the initial model (Figure 4A). Furthermore, RMSD distributions converged towards a common dominant population (Figure 4B), indicating that the assembly relaxes towards a similar structural arrangement and remains stable on the microsecond timescale. Representative CG conformations were subsequently backmapped to atomistic resolution and used as starting structures for independent 200 ns all-atom (AA) molecular dynamics simulations (Supp. Figure 10B). In all simulations, the two monomers remained associated throughout the trajectories, indicating that the dimeric assembly identified during the CG simulations is preserved at atomic resolution.

However, analysis of the intersubunit centre-of-mass (COM) distance revealed two distinct structural behaviours (Figure 4C). Replicates 1 and 2 converged towards a common compact arrangement, whereas replicate 3 progressively sampled a more extended configuration with increased intersubunit separation. This distinction was further supported by RMSD distributions, which identified two predominant conformational populations corresponding to these states (Figure 4D,E).

Representative structures corresponding to the closed and open arrangements were further evaluated using PDBe PISA, PRODIGY and PRODIGY Crystal (Supp. Table 3). These analyses consistently identified only the compact arrangement as a biologically relevant dimeric assembly. Accordingly, this interface displayed a substantial buried surface area (2254 Å²), favourable interaction energetics (ΔG_int_ = −9.4 kcal mol^−1^), and a strong predicted binding affinity (K_d_ = 4.9 nM), while PDBe PISA (CSS = 1.0) and PRODIGY Crystal (probability = 0.61) both supported its biological relevance. The interface is further characterized by a predominance of hydrophobic contacts and by the presence of a pronounced electronegative cleft extending across the dimer interface. In contrast, the extended arrangement failed to meet these criteria, suggesting that it most likely represents a partially destabilized conformation during the simulation rather than an alternative biologically relevant dimeric state (Supp. Table 2).

Structural comparison between the initial AF3 model and the biologically relevant dimeric assembly revealed a marked reorientation of the conserved C227–C228 vicinal disulfides while preserving their overall location within the dimer interface. In the initial AF3 model, the disulfides adopt an inward-facing orientation within the dimer interface, a region displaying slightly reduced local confidence (pLDDT ∼80) (Supp. Figure 9A). During the AA simulations, partial loss of helicity at the C-terminal end of the α-helix containing the C227–C228 pair enables reorientation of the disulfides. As a result, the opposing disulfides adopt a face-to-face arrangement across the dimer interface, becoming solvent-exposed (Figure 4F).

Given the proposed role of extracellular cysteine-containing motifs in ROS-responsive signaling by plant receptor kinases (Wu et al., 2020; Sun et al., 2025), particular attention was given to the structural environment of the C227-C228 region.

To explore whether these structural features are accompanied by distinctive chemical properties, pKa values were estimated for all cysteine residues using PROPKA. Because pKa calculations require free thiol groups, the analysis was performed on the reduced form of the cysteine residues. Whereas most cysteines of the ECD displayed predicted pKa values between 8.4 and 16.0, C227 and C228 were clear outliers, with pKa values of 6.7-7.2 and 5.3-6.1, respectively. Similar values were obtained for the isolated monomer, indicating that these unusually low pKa values arise primarily from the local physicochemical environment of the C227-C228 pair rather than from dimerization itself. Analysis of the individual PROPKA energy contributions identified the local electrostatic environment, including favorable interactions with the adjacent R229 residue, as a major determinant of these pKa shifts (Supp. Table 4).

Together, these simulations indicate that the CRK18 ECD forms a stable homodimer that brings the C227-C228 vicinal disulfides into a closely apposed and solvent-exposed configuration. Analysis of the reduced cysteine pair further identifies C227 and C228 as the cysteine residues with the lowest predicted pKa values within the ECD, suggesting that their chemical environment may favour thiol deprotonation and be compatible with a potential role in redox-related regulation. Notably, the closed dimeric arrangement also generates a pronounced electronegative cleft at the dimer interface (Supp. Figure 11), which may represent a structurally accessible region potentially involved in ligand recognition or molecular interactions.

### CRK18 forms cell wall integrity-related protein complexes and regulates the nematode infection process

In accordance with the previous paragraph, we identified potential interactors of CRK18 that may serve as indicators of the signalling pathways in which CRK18 participates. To do that, we performed immunoprecipitation-mass spectrometry (IP-MS) on rosettes from two four-week-old Arabidopsis CRK18 overexpression (OE) lines: *35S:CRK18-3xFLAG-mScI* #1 (CRK18-OE1) and *35S:CRK18-3xFLAG-mScI* #2 (CRK18-OE2). CRK18-3xFLAG-mScI was immunoprecipitated using anti-FLAG beads from CRK18-OE1 and CRK18-OE2, with the background line (Col-0 roGFP2-Orp1) as a control.

We identified 23 proteins significantly enriched in CRK18-OE1 and 54 in CRK18-OE2 (p-value < 0.01, log2(Fold Change) >1) (Figure 5A, Supp Figure 12, Supp. Table 5). Gene Ontology (GO) analysis showed enrichment for proteins involved in defence response regulation, proton transfer, and pH homeostasis (Supp. Figure 12C). 23 proteins were co-immunoprecipitating with CRK18 in both lines, suggesting a relatively strong association and high technical reproducibility. Of the 23 proteins found in both lines, 17 localise to the plasma membrane (Figure 5A, Supp. Table 5). Notably, eight of the identified plasma membrane proteins were receptor kinases (RKs) (excluding CRK18), and eight others were pumps or channels (Figure 5A).

Four of the eight RKs identified in both lines have potential carbohydrate-binding domains (e.g., Lectin-like and malectin-like domains) in their extracellular regions. Based on the performed IP-MS on two independent lines, we identified Wall-Associated Receptor Kinase 1 (WAK1), Stress-Induced Factor 3 (SIF3), L-type Lectin Receptor-like Kinase IV.1 (LECRK-IV.1), and LETUM1 (LET1). Lectin domains are known to bind carbohydrates, for example, components from the cell wall(Molina et al. 2024). However, the lectin-like domain-containing RKs have also been shown to bind non-carbohydrate compounds, such as peptides and ATP(Chen et al. 2017; Stegmann et al. 2017). In addition to the lectin-like domain-containing proteins, we identified Proline-Rich Extensin-Like Receptor-like kinase 15 (PERK15), which has been proposed to bind the cell wall components (Invernizzi et al. 2022a). Three other RKs identified in both lines are MDIS1-interacting receptor-like kinase 2 (MIK2), Suppressor of BIR1 (SOBIR1), and Transmembrane kinase 4/BAK1-associating Receptor-like Kinase 1 (TMK4/BARK1). MIK2 and SOBIR1 are involved in immune responses, while TMK4/BARK1 may regulate Auxin biosynthesis(Dai et al. 2013; Gust and Felix 2014; Albert et al. 2015; Hou et al. 2021; Rhodes et al. 2021).

In addition to RKs, we identified a number of plasma membrane pumps in both OE lines, including four autoinhibited plasma membrane proton ATPases (AHAs) (AHA1, AHA2, AHA3, AHA11), three ATP-binding cassette (ABC) transporter G family members (ABCG) (ABCG12, ABCG22, ABCG36), and Sucrose transport protein 2 (SUC2). These proteins are involved in transport, in maintaining cellular homeostasis and stress responses, such as modulating apoplastic pH and exporting antimicrobial compounds(Elmore and Coaker 2011; Gräfe and Schmitt 2021).

We also examined potential changes in the proteome and phosphoproteome levels in rosettes from four-week-old Arabidopsis of two independent CRK18 overexpression lines (CRK18-OE1 and CRK18-OE2) of the Col-0 ecotype and compared them to the control (Supp. Tables 6 and 6). We detected a few significant changes in the abundance or phosphorylation of proteins involved in cell-surface signalling. This lack of minimal changes could indicate that CRK18 requires a trigger for activation and possibly complex reorganisation leading to activation of specific signalling pathways. Interestingly, CRK18 is constitutively phosphorylated on Tyrosine 498 (Tyr 498), suggesting this phosphosite is required for the complex’s organisation with identified interactors.

Receptor kinases operate as dynamic interaction networks within the PM. The PM is dynamically organized into various nano-environments, and the initiation of signalling cascades has repeatedly been shown to correlate with changes in the nanoscale dynamics of signalling modules(Arx et al. 2026a). For instance, BAK1 is spatially arrest to pre-formed FLS2 nanodomains upon flg22 perception (Arx et al. 2026a). Similarly, BIK1 is dynamically stabilized within the plasma membrane upon flg22 perception(Wang et al. 2024). Therefore, we aimed to analyse the nanoscale organisation of CRK18 at the plasma membrane. To this end, we used photochromic reversion, a variant of single-particle tracking photoactivated localisation microscopy (spt-PALM) that enables long-term single-molecule imaging (Arx et al. 2024). We used CRK18 tagged with the photoconvertible fluorescence protein mEOS3.2 expressed under a constitutive promoter in *N. benthamiana*. Within minutes following flg22 treatments, CRK18-mEOS3.2 exhibits a decreased lateral diffusion and a lower diffusion coefficient subsequent (Supp. Figure 13A and B). We used computational analysis of spatial arrests (CASTA) to analyse dynamic changes in CRK18 diffusion. Flg22 treatment led to global changes in CRK18-mESO3.2 diffusion associated with an increase in the duration CRK18-mEOS3.2 spent in a spatial arrested state, without affecting the area or the frequency of arrest events (Supp. Figure 13 C and D). This evidence, along with the identification of immune-related proteins, including various receptor kinases with lectin-like domains, as interacting partners, suggests that CRK18 may play a regulatory role in plant stress responses.

CRK18, but not CRK16 or CRK17, two close homologs of CRK18, is highly transcriptionally upregulated upon nematode *Heterodera schachtii* infection. One of the identified interacting partners of CRK18 is PERK15, and this family of proteins is generally thought to be putative sensors of cell wall integrity, linking cell wall status and mechanical cues to intracellular signalling. Therefore, to investigate the *in vivo* role of CRK18 during stress responses induced by nematode parasitism, we performed infection assays using the beet cyst nematode *Heterodera schachtii*. Arabidopsis mutant lines, including two independent *crk16/17/18* triple knockout lines and a CRK18 overexpression line (CRK18-OE2), were inoculated with infective second-stage cyst nematodes. The progression of the nematode life cycle was evaluated at 14 and 28 days post-inoculation (dpi) (Figure 5B, C). At 14 dpi, early infection parameters were assessed by quantifying the number of developing third-stage juveniles (J3s), young cysts, and the total nematode count. Our analysis revealed that both *crk16/17/18* triple mutant lines exhibited a significantly higher number of total infections than wild-type controls, indicating that the absence of these CRKs enhances plant susceptibility to cyst nematodes. Conversely, the CRK18 overexpression plants displayed a marked reduction in overall infection levels at this early nematode developmental stage (Figure 5B, Supp. Figure 14A, B). To determine the impact of these genetic alterations on nematode development and reproduction, we monitored the infection at 28 dpi. At this later time point, we recorded the number of fully developed cysts, J3s, adult males, and the total nematode count. Consistent with our 14 dpi observations, the triple *crk16/17/18* knockout mutants showed significantly higher nematode numbers, indicating that these CRKs are part of a coordinated defense response restricting nematode parasitism (Figure 5C, Supp. Figure 14C-E). In contrast, the CRK18 overexpression mutant showed reduced susceptibility at 28 dpi. Taken together, these results suggest that CRK18 may act as a negative regulator of *H. schachtii* infection.

## Discussion

This study enhances understanding of CRK biology by investigating the biochemical and biophysical properties of CRK18 and integrating structural predictions with *in vitro* and *in planta* data. The combined evidence positions CRK18 as a membrane-localized, homodimerizing receptor-like kinase, residing in proximity with multiple RKs and whose ECD cysteines play distinct roles in folding and stability versus potential regulatory function. Furthermore, CRK18 association with plasma membrane receptors and transporters suggests that CRK18 is embedded within signalling modules that control cell wall–associated perception, apoplastic pH, and immune responses.

### Disulphide bonds maintain the stability of the CRK18 ECD and facilitate proper receptor trafficking

Cysteines, despite being among the least abundant yet most conserved amino acid residues in proteins, play roles in their structure, metal binding, catalysis, and redox chemistry. Our *in vitro* analyses highlight that the conserved cysteines in the DUF26 domains are central to CRK18-ECD stability. Mutation of the predicted disulphide bonds C216–C241 (DUF26B), C227–C228 (DUF26B, vicinal cysteines) and especially C94–C122 (DUF26A) substantially reduced the thermal stability of the ECD (Figure 1A-D). The mutant variant CRK18-ECD^C94A,C122A^ exhibits a broad unfolding transition and an estimated Tm of ∼30 °C, indicating pronounced destabilisation. This is consistent with structural predictions and previous work on DUF26-containing proteins (Plasmodesmata Localized Proteins, PDLPs), where conserved cysteine networks are thought to stabilise the β-sandwich fold and maintain the relative orientation of the two DUF26 modules(Vaattovaara et al. 2019). The modest increase in Tm at lower pH observed in wild-type CRK18-ECD, and even more pronounced in some cysteine mutants, may reflect the mildly acidic apoplastic environment and suggests that apoplastic pH helps maintain the structural integrity of poorly folded or partially destabilised CRK18 variants.

Consistent with the biochemical *in vitro* data, most cysteine-to-alanine mutants in the full-length receptor were mislocalised or partially retained in the ER (Figure 2). These observations suggest that correct disulphide bond formation in the ECD is required for folding, efficient ER quality-control passage, and delivery to the plasma membrane, similarly to what is known for other receptor-like kinases and secretory proteins(Brast et al. 2011; Hurst et al. 2019). The one notable exception was the C227A/C228A mutant, which retained robust plasma membrane localisation and showed only a modest decrease in ECD stability, indicating that the vicinal cysteines are not essential for folding or trafficking but may play a more regulatory role. These cysteines are found in some other members of the CRK family from the variable clade, but not in those from the basal clade or the previously mentioned PDLPs, which are similar to the CRK basal clade (Vaattovaara et al. 2019; Martin-Ramirez et al. 2025; Stouthamer et al. 2026). This could imply functional diversification between different DUF26-containing proteins.

### Limited impact of redox perturbation and vicinal cysteines on CRK18 homodimerization

Work by Martin Ramirez et al. demonstrated that CRK18-ECD homodimerization depended on ROS, leading to dissociation of the homodimeric complex upon ROS treatment (Martin-Ramirez et al. 2026). We therefore further tested whether extracellular redox changes and specific cysteines directly affect CRK18 homodimerization. Using SEC-MALS and mass photometry, we confirmed that CRK18-ECD forms monomer–dimer species *in vitro* (Figure 3A, Supp. Figure 1A). MST measurements revealed that CRK18-ECD homodimerizes with affinities in the low to mid-nanomolar range at both pH 5 and 7.5, with only modest pH dependence (Figure 3 B, C). This is a high-affinity interaction, as other RK homodimers described in the literature display affinities in the sub-micromolar to low micromolar range(Liu et al. 2012). Moreover, neither oxidative (H_2_O_2_) nor reductive (DTT) treatment within the tested range had a strong effect on dimerization affinity, despite DTT clearly compromising ECD thermal stability at higher concentrations (Supp. Figure 7). Together, these data suggest that the disulphide network is important for overall structural integrity but that the dimerization interface itself is relatively robust to moderate redox perturbations, at least in the absence of additional cellular factors.

Importantly, the vicinal cysteines C227 and C228 were dispensable for homodimerization of both the ECD *in vitro* and full-length CRK18 *in planta*. The purified CRK18^C227A, C228A^-ECD mutant dimerized with almost the same affinity as wild-type at pH 7.5 and the full-length showed no detectable difference in Co-IP experiments (Figure 3D, E). Thus, homodimer formation appears to be a constitutive property of CRK18 that does not require the vicinal disulphide.

Multiscale molecular dynamics simulations indicate that the CRK18 ECD forms a stable homodimer (Figure 4). Structural analyses identified a single biologically relevant dimeric assembly characterized by an extensive buried interface, favourable interaction energetics and a strong predicted binding affinity. A key structural feature of this assembly is the reorganization and solvent exposure of the C227-C228 vicinal disulfides. Although the initial AF3 model already positions these residues at the dimer interface, they are oriented towards the interior of the interface. During atomistic simulations, increased flexibility at the C-terminal region of the α-helix containing the C227-C228 pair enables local rearrangement and exposure of these residues. pKa calculations further predicted that C227 and particularly C228 display the lowest pKa values among CRK18 cysteines (Supp. Table 4). This tendency was largely preserved in the isolated monomer, indicating that their chemical properties arise primarily from the local environment of the cysteine pair rather than dimerization itself. Together, these findings identify a CRK18 dimeric assembly in which the C227-C228 pair displays structural and physicochemical features compatible with a potential role in redox regulation. Interestingly, this vicinal disulfide is not conserved across the CRK family, suggesting that this configuration may represent a specialized feature of CRK18 rather than a general characteristic of CRK ECDs.

Vicinal disulphides are well established as dynamic structural elements that regulate protein function by modulating local conformation and molecular interactions rather than global folding (Richardson 1981; De Araujo et al. 2013; Richardson et al. 2017b). In this context, the structural rearrangement identified here places the CRK18 C227–C228 pair in a solvent-exposed environment compatible with redox-dependent modification, although direct ROS sensing remains to be demonstrated. Similar extracellular cysteine-based regulatory mechanisms have recently been described for plant receptor kinases such as HYDROGEN-PEROXIDE-INDUCED CALCIUM INCREASES 1 (HPCA1; also known as CARD1: CANNOT RESPOND TO DMBQ), supporting the idea that local redox chemistry may represent a broader mechanism for regulating receptor activity(Laohavisit et al. 2020; Wu et al. 2020; Ishihama et al. 2026). Interestingly, vicinal disulphides have also been implicated in the organization of ligand-binding sites in other proteins, including glycan-binding pockets such as that described for Impaired in Glycan Perception 1 (IGP1, also known as Cellooligomer Receptor Kinase 1 (CORK1)) (Jiménez-Sandoval et al.). The C227–C228 pair in CRK18 generates a pronounced electronegative cleft at the dimer interface (Supp. Figure 11). This observation raises the possibility that CRK18 dimerization may contribute to the formation of an extracellular recognition surface, providing a structural framework for future investigation of potential ligand interactions.

### CRK18 associates with receptors and transporters that regulate stress responses and apoplastic homeostasis

To place the role of CRK18 in the signalling context, we identified interacting proteins in Arabidopsis lines overexpressing CRK18. The IP-MS experiments revealed a robust set of 23 proteins identified from two independent lines, with the majority localising to the plasma membrane (Supp. Figure 9). Notably, we found eight receptor kinases and multiple transporters and pumps. This, together with the fact that, based on the phosphoproteome results, the CRK kinase domain remains constitutively phosphorylated on Tyr 498, suggests that CRK18 is part of a broader cell-surface signalling and transport hub rather than an isolated receptor.

Among the receptor kinases, several contain lectin- or malectin-like extracellular domains (e.g. WAK1, SIF3, LECRK-IV.1, LET1), suggesting that CRK18 may be functionally linked to receptors involved in cell wall or carbohydrate-related perception, but potentially also non-carbohydrate ligands, such as peptides or extracellular nucleotides. This observation is particularly interesting because members of the DUF26 protein family have also been linked to glycan recognition. The single DUF26-containing Cysteine-Rich Repeat Secretory Protein (CRRSP) Ginkbilobin2 (GNK2) from *Ginkgo biloba* exhibits antifungal activity, and the NMR structure of the GNK2-mannose complex reveals a mannose-binding motif composed of three residues: N11, R93, and E104 (Miyakawa et al. 2014). Similarly, two CRRSPs containing two DUF26 domains from maize, AFP1 and AFP2, have also been shown to bind mannose(Ma et al. 2018). Additionally, wheat’s TaCRK3 displays antifungal activity and contains conserved mannose-binding residues(Guo et al. 2021). Several Arabidopsis CRKs contain the residues corresponding to the GNK2 mannose-binding motif site, but CRK18 lacks this motif (Stouthamer et al. 2026). Nevertheless, the presence or absence of the GNK2 residues is not necessarily predictive of mannose or other glycan perception. Recently, CRK7 was identified as a receptor for wall teichoic acid(Pierdzig et al.). Although CRK7 contains the three residues corresponding to the GNK2 mannose-binding site, it does not contain all residues that contribute to mannose binding in GNK2. Together, these findings suggest that CRKs may recognize mannose or other glycans, potentially through an interaction interface distinct from that of GNK2. Additional interactors, such as PERK15, further underscore a possible role in sensing or transmitting information across the cell wall–plasma membrane continuum. PERK15 belongs to a small family of 15 receptor-like kinases that feature a proline-rich extracellular domain, resembling extensin cell wall proteins, and they are thought to function as cell wall integrity sensors often related to nematode infection processes (Bohlmann and Sobczak 2014; Invernizzi et al. 2022b; Pijnacker et al. 2026). Transcriptional upregulation of CRK18 during infection by the nematode *Heterodera schachtii* reported by Pijnacker et al. 2026, together with higher susceptibility, as indicated by higher nematode numbers in the crk16/17/18 knockout lines and conversely lower nematode numbers in the CRK18-OE2 lines, clearly suggests its negative regulatory role during the infection process. This underlines the importance of CRK18 in responding to changes in cell wall integrity. Additionally, the enrichment of plasma membrane H^+^-ATPases (AHA1/2/3/11), ABCG transporters, and SUC2 indicates that CRK18 might influence, or respond to, apoplastic pH and transport processes(Elmore et al. 2021). Proton pumps and ABC transporters are key regulators of cell wall extensibility, membrane potential, and the export of antimicrobial metabolites, all of which are central to growth and stress responses. The observation that apoplastic pH modulates CRK18-ECD stability, together with the association of CRK18 with proteins involved in proton transfer and pH homeostasis, suggests that CRK18 participates in feedback loops linking cell-surface perception to apoplastic and membrane transport dynamics.

We also identified MIK2 and SOBIR1, which are established components of plant immune receptor complexes(Wu et al. 2024; Zhai et al. 2024). Consistent with a role in stress and immunity, super-resolution imaging revealed that CRK18 mobility at the plasma membrane decreases after flg22 elicitation, a well-known bacterial PAMP, and our interaction data include multiple immune-related receptor kinases (Supp. Figure 13). The co-receptor BAK1 and the receptor-like cytoplasmic kinase BIK1 are known to reorganize into nanoclusters upon ligand perception (Wang et al. 2024; Arx et al. 2026a). The changes in CRK18 mobility behaviour after flg22 treatment suggest that CRK18 may be recruited into, or rearranged within, such signalling nanodomains. Although our phosphoproteomics did not reveal extensive rewiring of cell-surface signalling components under the tested conditions, CRK18 itself is constitutively phosphorylated on Tyr 498, suggesting potential regulation of CRK18 activity or interaction capacity via tyrosine phosphorylation (Supp.Table 7).

In conclusion, our work shows that CRK18 is a structurally constrained, disulphide-stabilised receptor that homodimerizes at the plasma membrane and is in proximity with receptor kinases and transporters involved in stress signalling and apoplastic homeostasis. The conserved cysteines ensure proper folding and localization, whereas the solvent-exposed vicinal disulphide emerges as a promising candidate for a redox-regulated switch that may tune CRK18 function during plant responses to environmental challenges.

### A working model for CRK18 function and regulation

Integrating our findings, we propose the following model where CRK18 forms stable, disulphide-stabilised homodimers at the plasma membrane, with conserved DUF26 cysteines required for proper folding and ER export (Figure 6). In this basal state, CRK18 associates with receptors and transporters, including lectin-domain receptor kinases, immune receptors, and pumps/transporters controlling apoplastic pH and metabolite fluxes. The vicinal C227–C228 disulphide is not necessary for this basal organisation but represents a solvent-exposed, structurally permissive redox-sensitive element that may fine-tune CRK18 function under specific oxidative or stress conditions, for example, by locally altering the ECD conformation to allow for ligand binding or modulating affinity for particular co-receptors or ligands. Changes in apoplastic pH, redox status, or cell wall integrity, as for instance occurring during pathogen attack or abiotic stress, may then be sensed or integrated by CRK18 in conjunction with its associated receptors and transporters, leading to changes in receptor mobility, complex composition, and downstream signalling.

### Future directions

While our study establishes foundational principles of CRK18 biochemistry and potential physiological roles, it also opens novel directions linking CRK to cell wall signalling. Future work should therefore focus on identifying physiological ligands or cell wall–derived cues that modulate CRK18 activity, ideally in the context of endogenous expression. Dissecting the functional relevance of the vicinal C227–C228 motif will require targeted redox-proteomics, site-specific redox reporters, or genetically encoded sensors to monitor its redox state during stress responses. Genetic analyses of *crk18* loss-of-function and overexpression lines, in combination with mutants of key interactors (e.g., PERK15, WAK1, MIK2, SOBIR1, AHAs, and ABCGs), will be important for placing CRK18 within defined immune and pH-regulatory pathways. Finally, structural studies of CRK18-ECD, alone and in complex with candidate ligands or co-receptors, could clarify how the conserved disulphide network and the vicinal cysteines shape receptor conformation and interaction interfaces.

### Author contribution

E.S-L, J.S and J.L designed the study. J.S and J.L performed protein expression, purification and biochemical characterization with the help of S.L. G.V-P, A.M and M.G-A performed structural predictions. D. Ž and J.B performed the mass photometry assay. J.M, S.R and A.O were involved in FRET-FLIM measurement. J.S. performed localization studies. J. S and J.L, with the help of M.R and S.B performed all (phosphor) proteomics and IP-MS. M.v.O and C.G shared insect cell lines and provided expertise on maintaining the cultures. M.v.A and J.G performed the nanoscale organization assay. R.S.L and J.L.T performed the nematode infection assay. E.S-L, J.S and J.L wrote the paper.

## Acknowledgment

Measurements on the Prometheus Panta and Monolith (NanoTemper) were performed at the Protein Research Centre of Utrecht University. We would like to thank Prof. Dr Markus Schwarzländer from the Institute of Plant Biology and Biotechnology (IBBP), University of Münster, for kindly sharing the roGFP2-Orp1 line with us. We thank Dr Carlo van Mierlo for his input and feedback on the biophysical characterization of CRK41-ECD stability. We would also like to thank Dr Jan Willem Borst for providing the ER-Marker gene AtVMA21a.

## Supplementary material

Supplementary data in the form of figures are attached as a separate Word document. Structural predictions and proteomics data are available as tables.

## Funding

This work was supported by grants PID2024-159175OB-I00 and CEX2020-000999-S (to AM) funded by Spanish Ministry of Science, Innovation and Universities (MCIU/AEI/10.13039/501100011033 and by “ERDF/EU”). GVP was supported by PRE2021-100446 fellowship funded by MCIU/AEI/10.13039/501100011033 and by FSE+. Research in the J.R.B. laboratory is supported by the Royal Society (URF\R1\211567) and UK Research and Innovation Frontier Research Guarantee Grant (EP/Y036158/1). D.Ž. is supported by the European Research Executive Agency under the HORIZON TMA Marie Skłodowska-Curie grant agreement (101205973). The work of E.S.-L. was supported by the NWO Talent Programme Vidi grant VI.Vidi.193.074

## Conflicts of interest

None declared.

## Data availability statement

Data are available in a repository and can be accessed using a unique identifier other than a DOI.

## Notes

### Competing Interest Statement

The authors have declared no competing interest.

