## Supplementary data file for "Structural and functional insights into the role of Cysteine-Rich Receptor-Like Kinase 18 (CRK18) in Arabidopsis"

### Supplementary data files

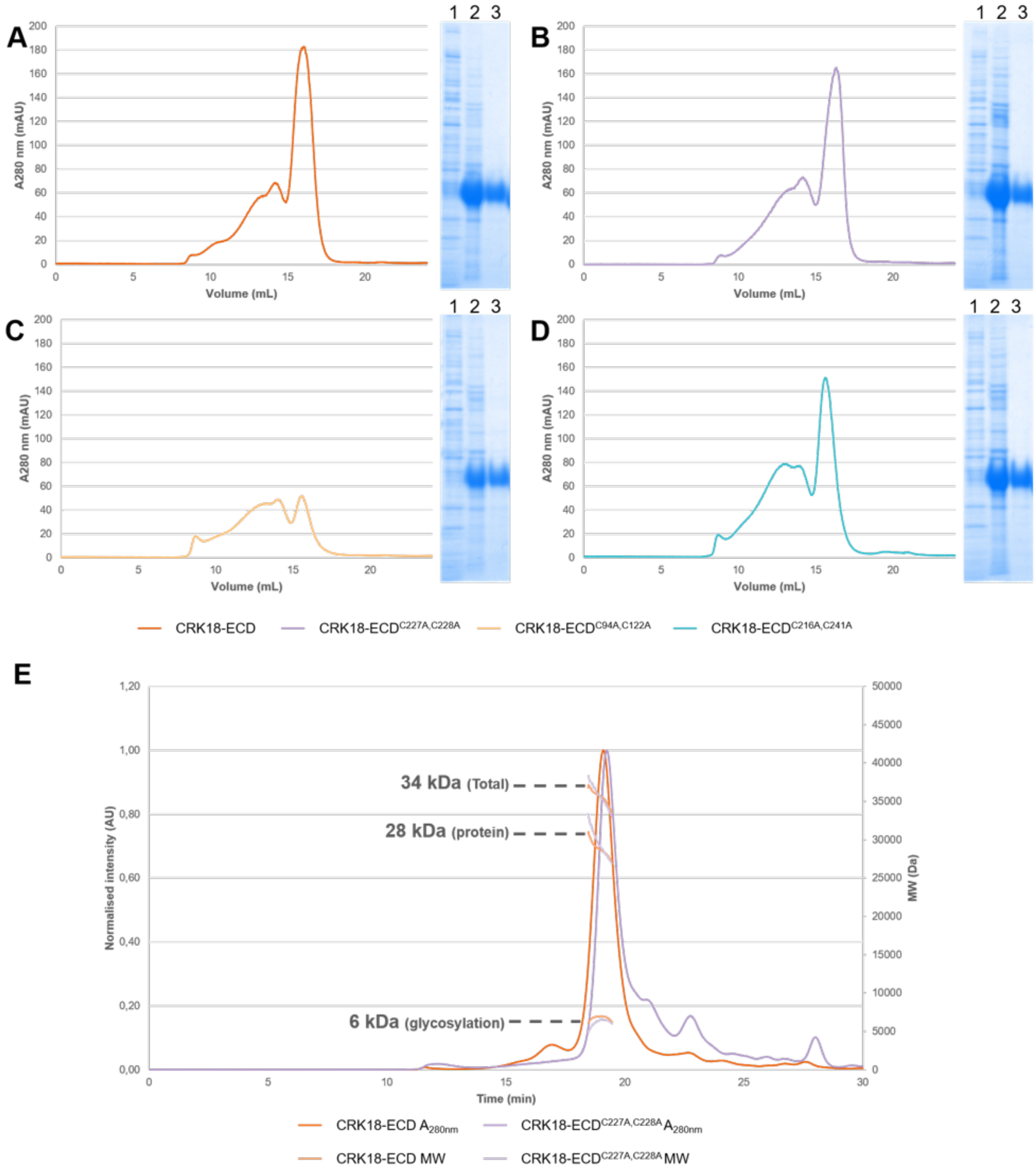


**Supplementary Figure 1.** Protein purification of CRK18-ECD and mutant variants. Size exclusion chromatography (SEC) runs and SDS-PAGE gels (from left to right: supernatant from Insect cells, pooled eluate from affinity chromatography, pooled eluate from SEC) for A) CRK18-ECD, B) CRK18-ECD$^{C227A,C228A}$, C) CRK18-ECD$^{C94A,C122A}$, D) CRK18-ECD$^{C216A,C241A}$. CRK18 and mutant variants started eluting ~15 mL, fractions from this main peak was pooled. E) SEC-MALS chromatogram of the purified CRK18-ECD. CRK18-ECD sample eluted from a Superdex 200 column as one peak. The CRK18-ECD protein size was determined as MW of 34kDa, of which 28 kDa was protein and 6 kDa was glycosylation, corresponding to a monomer of CRK18-ECD. SEC and SEC-MALS samples were in 20 mM HEPES, 150 mM NaCl, pH 7.5.

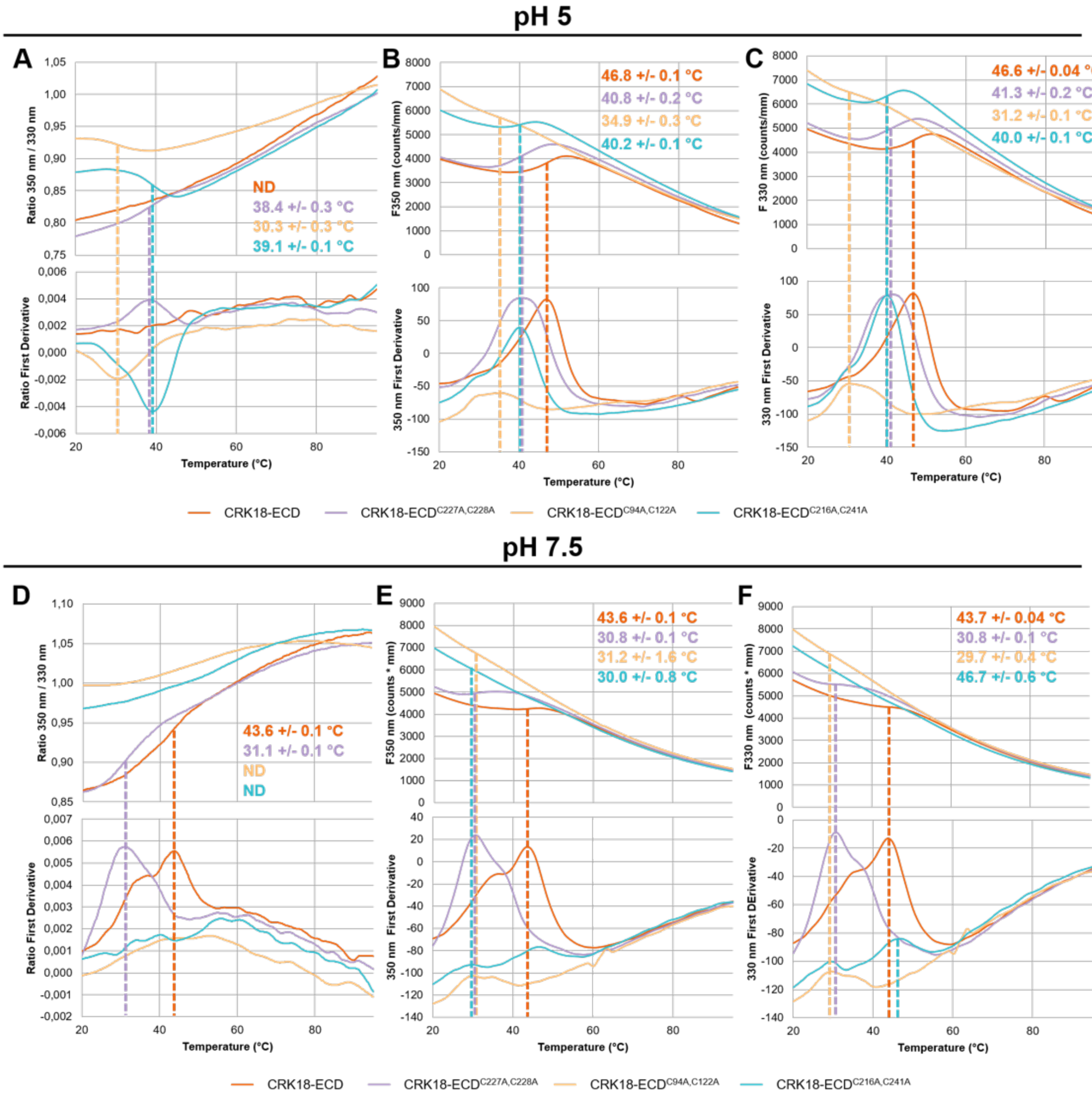


**Supplementary Figure 2. Thermal stability data of CRK18-ECD and mutant variants at pH 5 and pH 7.5.** nanoDSF measurements were performed for CRK18-ECD, CRK18-ECD$^{C227A,C228A}$, CRK18-ECD$^{C94A,C122A}$ and CRK18-ECD$^{C216A,C241A}$ at A-C) pH 5 or D-F) pH 7.5. A & D) Thermal unfolding curves showing the ratio of fluorescence emission at 350 and 330 nm and the first derivative of this ratio as a function of temperature. Fluorescence emission measured at B & E) 350 and C & F) 330 nm and the first derivative thereof as a function of temperature. From the temperature-dependent 350 and 330 nm fluorescence intensities and the ratio thereof the thermal midpoint ($T_m$) of the protein can be derived as the temperature at which 50% of the protein molecules is unfolded. All samples were prepared at a protein concentration of 0.4 mg/mL. pH 5 samples were prepared in 20 mM NaOAc, 150 mM NaCl, 0.1% Pluronic acid. pH 7.5 samples were prepared in 20 mM HEPES, 150 mM NaCl, 0.1% Pluronic acid.

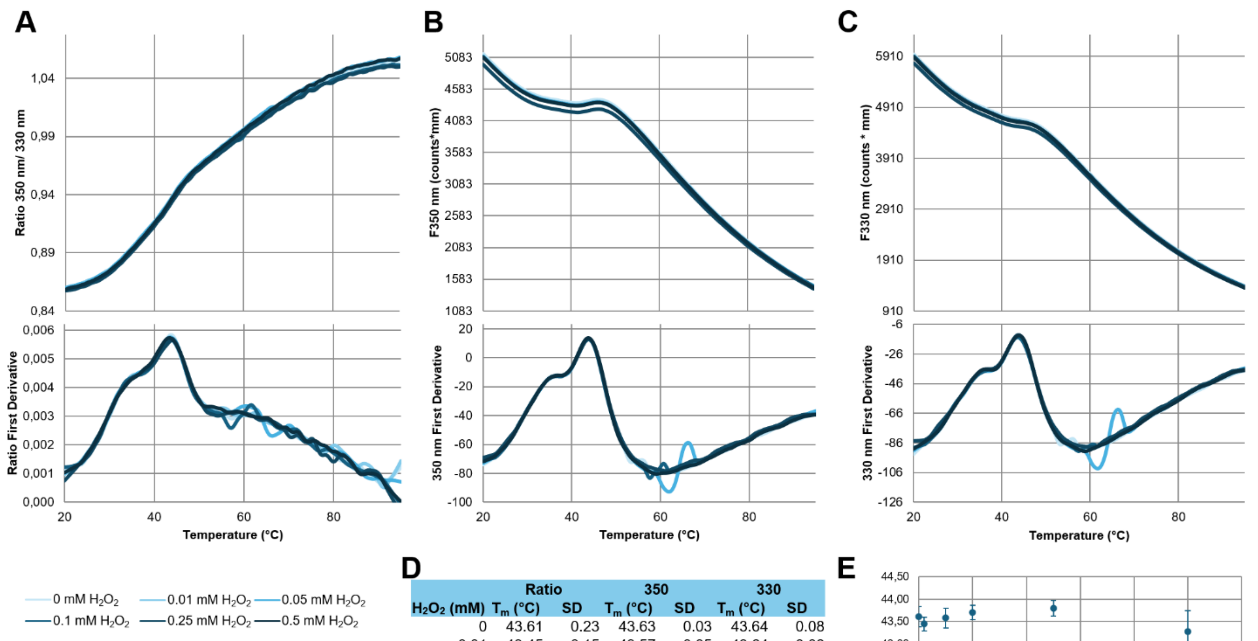


D

| $H_2O_2$ (mM) | Ratio $T_m$ (°C) | Ratio SD | 350 $T_m$ (°C) | 350 SD | 330 $T_m$ (°C) | 330 SD |
|---|---|---|---|---|---|---|
| 0 | 43.61 | 0.23 | 43.63 | 0.03 | 43.64 | 0.08 |
| 0.01 | 43.45 | 0.15 | 43.57 | 0.05 | 43.64 | 0.08 |
| 0.05 | 43.58 | 0.22 | 43.62 | 0.10 | 43.64 | 0.01 |
| 0.1 | 43.70 | 0.16 | 43.66 | 0.10 | 43.58 | 0.08 |
| 0.25 | 43.79 | 0.17 | 43.72 | 0.02 | 43.72 | 0.03 |
| 0.5 | 43.28 | 0.46 | 43.65 | 0.05 | 43.79 | 0.09 |

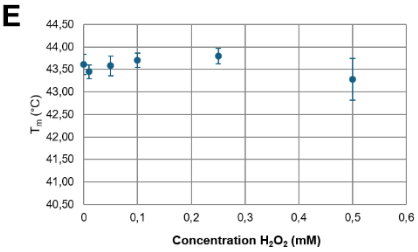


**Supplementary Figure 3. $H_2O_2$ does not affect the thermal stability of CRK18-ECD.** nanoDSF measurements were performed on CRK18-ECD at different $H_2O_2$ concentrations. All samples were prepared at a protein concentration of 0.4 mg/ml in 20 mM HEPES, 150 mM NaCl, 0.1% Pluronic acid pH 7.5, with varying concentrations of $H_2O_2$ (0-0.5 mM). A) Thermal unfolding curves showing the ratio of fluorescence emission at 350 and 330 nm and the first derivative of this ratio as a function of temperature. Fluorescence emission measured at B) 350 and C) 330 nm and the first derivative thereof as a function of temperature. D) From the temperature-dependent 350 and 330 nm fluorescence intensities and the ratio thereof the temperature at which 50% of the protein is unfolded (Tm) is determined. The table shows the Tm-values and corresponding standard deviations (SD). E) Tms of CRK18-ECD at different $H_2O_2$ concentrations.

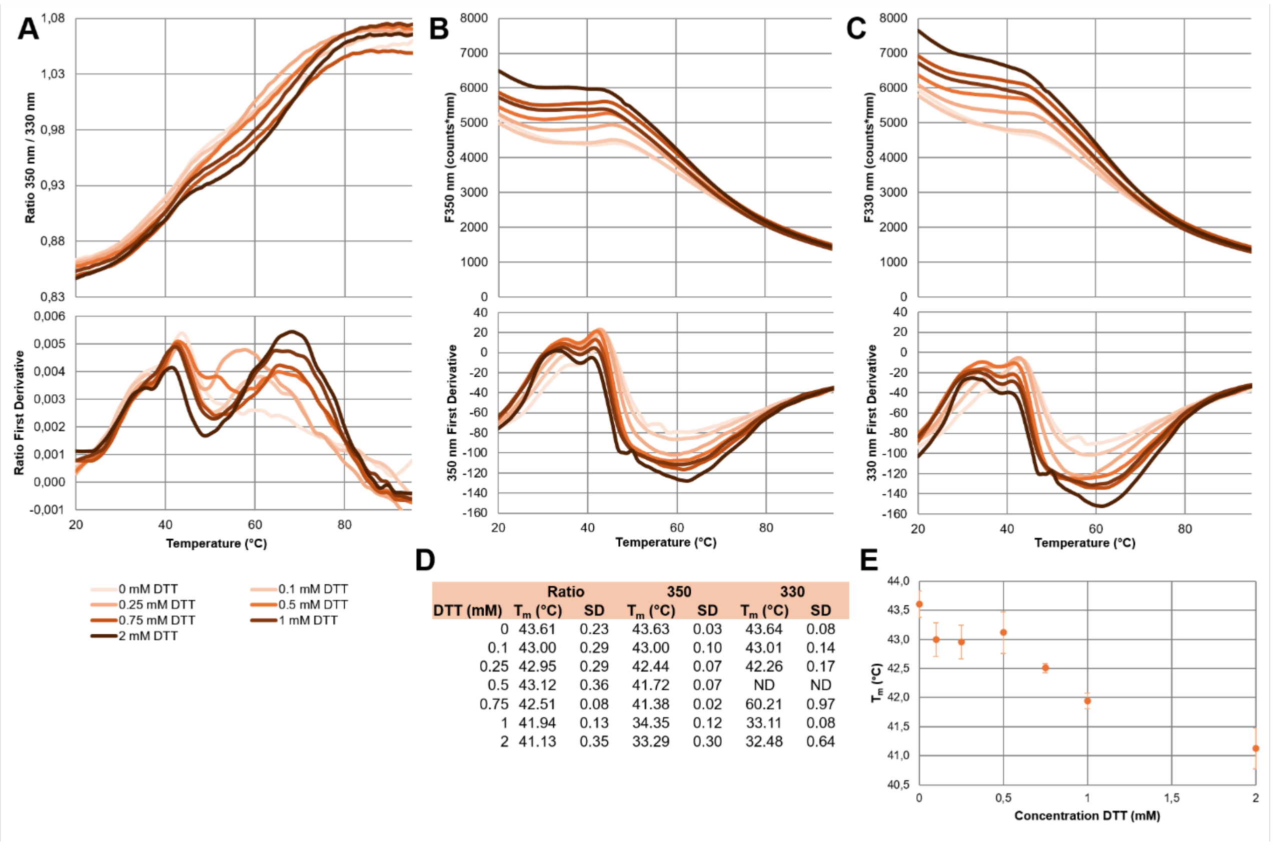


| | Ratio | | 350 | | 330 | |
|---|---|---|---|---|---|---|
| DTT (mM) | $T_m$ (°C) | SD | $T_m$ (°C) | SD | $T_m$ (°C) | SD |
| 0 | 43.61 | 0.23 | 43.63 | 0.03 | 43.64 | 0.08 |
| 0.1 | 43.00 | 0.29 | 43.00 | 0.10 | 43.01 | 0.14 |
| 0.25 | 42.95 | 0.29 | 42.44 | 0.07 | 42.26 | 0.17 |
| 0.5 | 43.12 | 0.36 | 41.72 | 0.07 | ND | ND |
| 0.75 | 42.51 | 0.08 | 41.38 | 0.02 | 60.21 | 0.97 |
| 1 | 41.94 | 0.13 | 34.35 | 0.12 | 33.11 | 0.08 |
| 2 | 41.13 | 0.35 | 33.29 | 0.30 | 32.48 | 0.64 |

**Supplementary Figure 4.** DTT affects the thermal stability of CRK18-ECD. nanoDSF measurements were performed on CRK18-ECD at different DTT concentrations. All samples were prepared at a protein concentration of 0.4 mg/ml in 20 mM HEPES, 150 mM NaCl, 0.1% Pluronic acid pH 7.5, at varying concentrations of DTT (0-2mM). A) Thermal unfolding curves showing the ratio of fluorescence emission at 350 and 330 nm and the first derivative of this ratio as a function of temperature. Fluorescence emission measured at B) 350 and C) 330 nm and the first derivative thereof as a function of temperature. D) From the temperature-dependent 350 and 330 nm fluorescence intensities and the ratio thereof the temperature at which 50% of the protein is unfolded (Tm) is determined. The table shows the Tm-values and the corresponding standard deviations (SD). E) Tms of CRK18-ECD at different DTT concentrations.

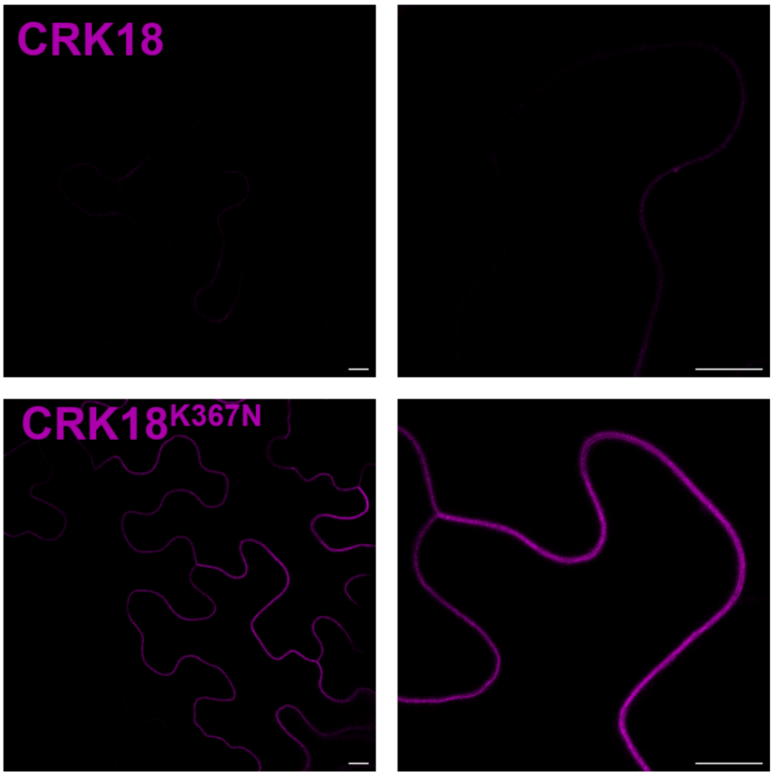


**Supplementary Figure 5. Inactivation of the kinase domain of CRK18 improves its transient expression in *N. benthamiana*.** 35S:CRK18-3xFLAG-mScI and 35S:CRK18$^{K367N}$-3xFLAG-mScI transiently expressed in epidermal cells of *N. benthamiana*. CRK18-mScI has low expression levels compared to CRK18$^{K367N}$-mScI. Scale bars are 10 µm.

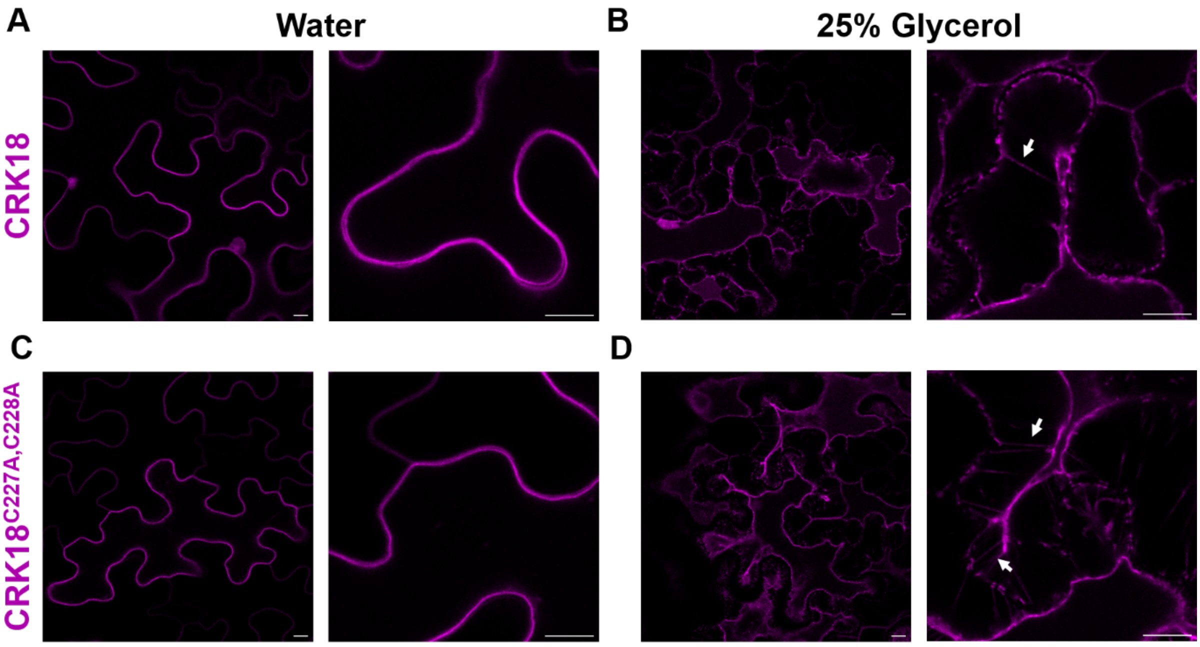


**Supplementary Figure 6. CRK18 and CRK18$^{C227A,C228A}$ localize to the plasma membrane.** A) 35S:CRK18-mScI expression in *N. benthamiana* leaf epidermal cells. B) Hechtian strands (white arrow) are visible after separation of the plasma membrane from the cell wall was induced by treatment with 25% glycerol, indicating localization of CRK18-mScI to the plasma membrane. C) CRK18$^{C227A,C228A}$-mScI shows a similar localisation as CRK18-mScI, and D) Hechtian strands (white arrows) are formed upon plasmolysis. Scale bars are 10 µm. All constructs contained the CRK18$^{K367N}$mutation to inactivate the kinase.

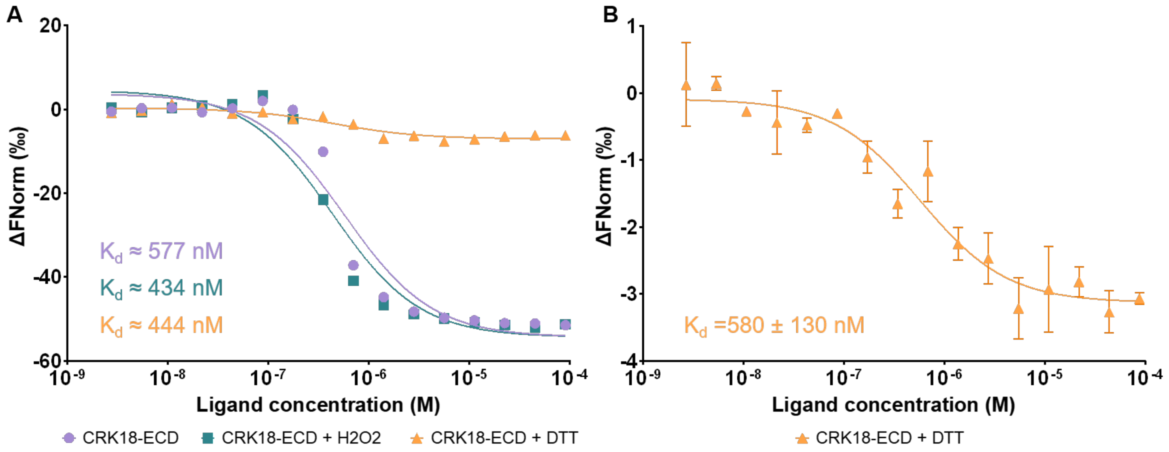


**Supplementary Figure 7. CRK18-ECD homodimerization is unaffected by pH, $H_2O_2$, DTT** A) Single MST measurements (n=1) of CRK18 homodimerization without treatment and treated with either 100 µM $H_2O_2$ or 0.5 mM DTT for one hour at 21 °C at pH 7.5 D) Dose-response curve of CRK18-ECD - CRK18-ECD measured at pH 7.5 after incubation with 0.5 mM DTT at 21 °C for one hour (n=3). For all MST measurements, a dilution series of unlabelled ligand was prepared, and labelled target was added at a constant concentration of 20 nM. The normalized change in fluorescence (ΔFNorm, ‰) was plotted against ligand concentration, from which the binding affinity (Kd ± SEM) could be determined. Samples were prepared in 20 mM HEPES, 150 mM NaCl pH 7.5, 0.1% Pluronic acid. MST infrared laser power was set to 40% for all experiments. Excitation power varied: (A) 100%, (B) 60%. Note that the Kd for CRK18-ECD homodimerization at pH 7.5 differs between measurements in Figure 3C) and Supp. Figure 7A), these measurements were performed on independent purifications of CRK18-ECD.

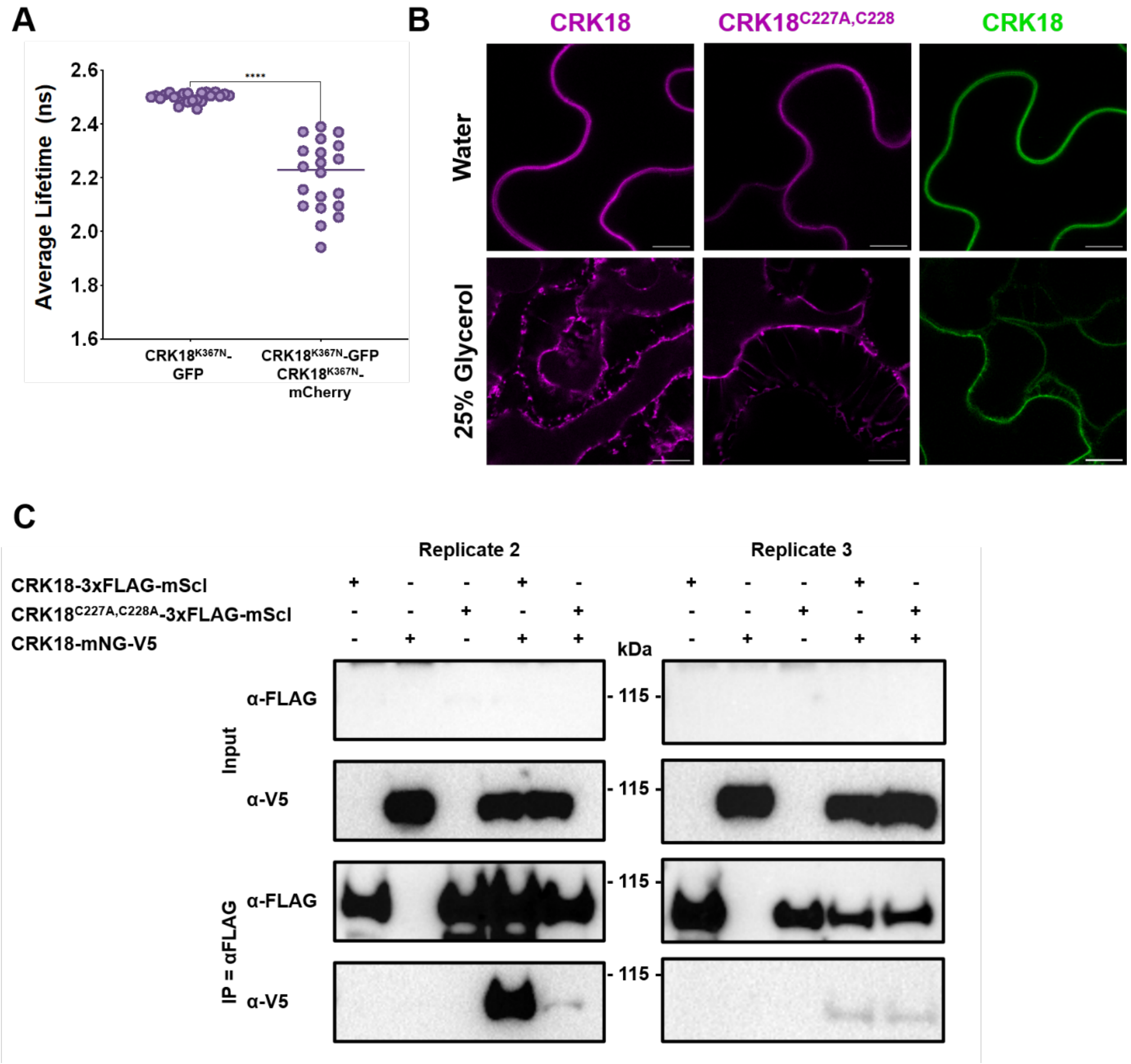


**Supplementary Figure 8. Replicates of Co-IP and FRET-FLIM experiments and supporting data.** A) FRET-FLIM analysis of kinase-inactive CRK18$^{K367N}$-GFP and CRK18$^{K367N}$-mCherry, showing a decrease in GFP lifetime upon co-expression, indicating dimerization. The average lifetime was quantified for 22 and 20 repeat measurements, respectively. Statistically significant difference was determined by Student's t-test, ****P<0.0001) B) Confocal microscopy images of CRK18-3xFLAG-mScI, CRK18$^{C227A,C228A}$-3xFLAG-mScI, and CRK18-mNG-V5 transiently expressed in *N. benthamiana*. After plasmolysis with 25% glycerol, Hechtian strands were observed, indicating the proteins were localised to the plasma membrane. Images were taken 2 dpi. Scale bars are 10 µm. C) Western blots showing replicates of Co-IPs experiments. In one of three repeats, CRK18$^{C227A,C228A}$ showed less protein compared to CRK18, while the other two repeats showed no difference.

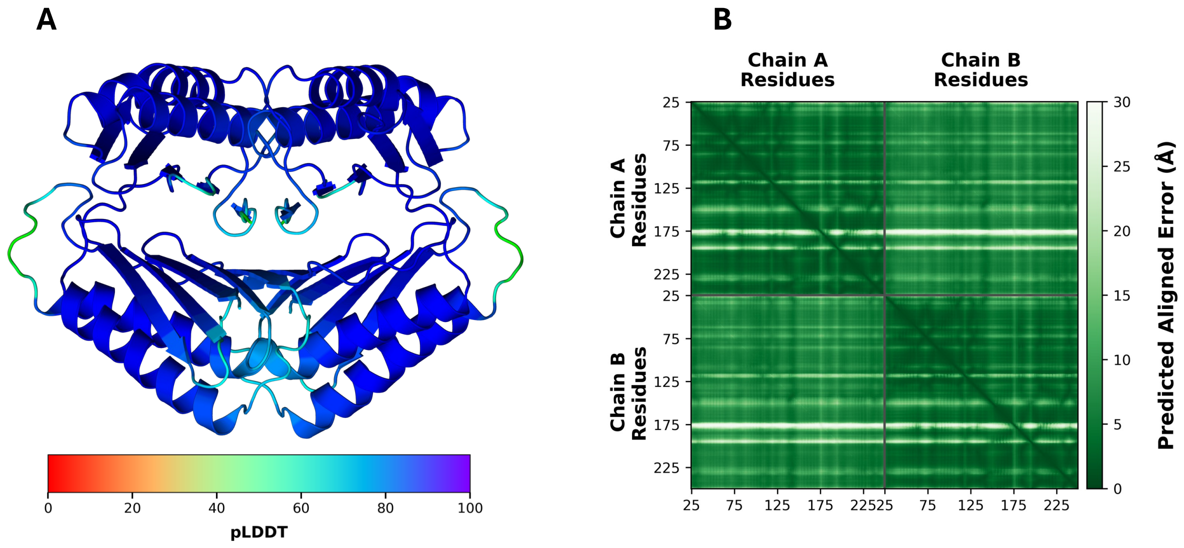


**Supplementary Figure 9. Confidence assessment of the AlphaFold3 prediction of the CRK18 ECD homodimer.** A) Structure colored according to pLDDT values. B) Predicted Aligned Error (PAE) matrix showing low alignment errors within and between monomers, supporting the predicted dimeric arrangement (ipTM = 0.76; pTM = 0.84). Overall, the model shows high local confidence (pLDDT > 90) across most of the structured core, with reduced confidence confined to flexible regions, including the N-terminal segment (residues 143-157), C-terminal segments (residues 189-198 and 225-234), and the loop spanning residues 171-184, where pLDDT values decrease below 80 and in some cases approach ~50, indicative of increased local flexibility.

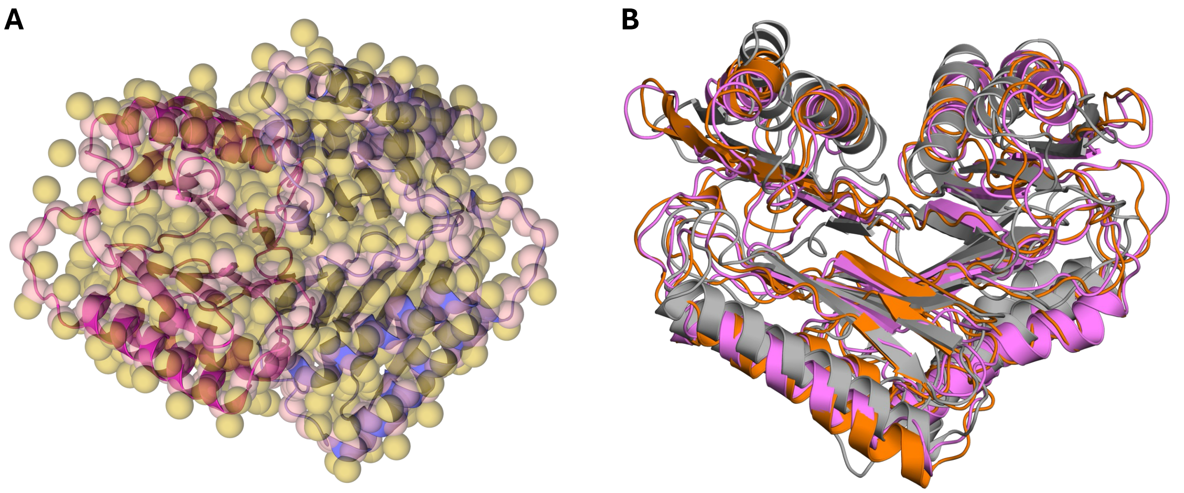


**Supplementary Figure 10. Coarse-grained representation of the CRK18 homodimer and structural convergence during CG simulations. A)** MARTINI CG model generated from the AF3 dimer used as the starting structure for all simulations. Monomers are shown in different colors. **B)** Structural superposition of the backmapped representative conformations extracted from the three independent CG simulations. Replicate 1 (pink) exhibits an RMSD of 1.33 Å relative to replicate 2 (orange) and 1.70 Å relative to replicate 3 (gray), while the RMSD between Replicates 2 and 3 is 2.10 Å.

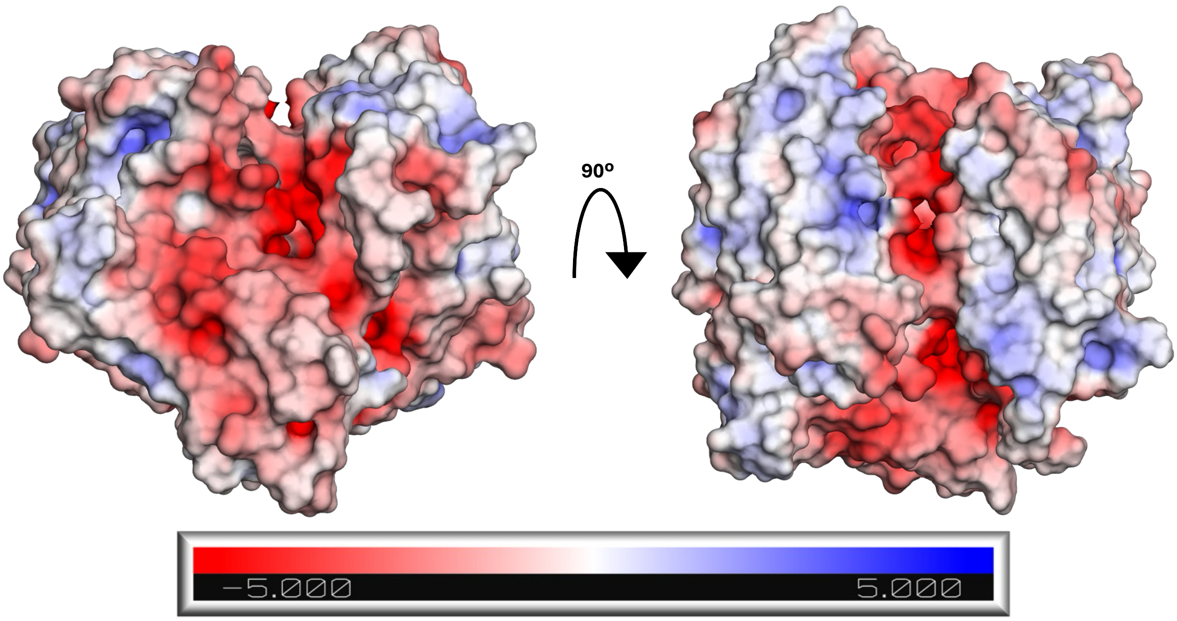


**Supplementary Figure 11. Electrostatic surface potential of the representative closed CRK18 ECD homodimer calculated by solving the Poisson–Boltzmann equation.** Two orthogonal views of the dimer are shown following a 90° rotation about the horizontal axis. The electrostatic surface reveals the formation of a pronounced negatively charged groove at the dimer interface, generated upon homodimerization. Electrostatic potential is shown from −5 to +5 kT/e, with negative and positive potentials colored red and blue, respectively.

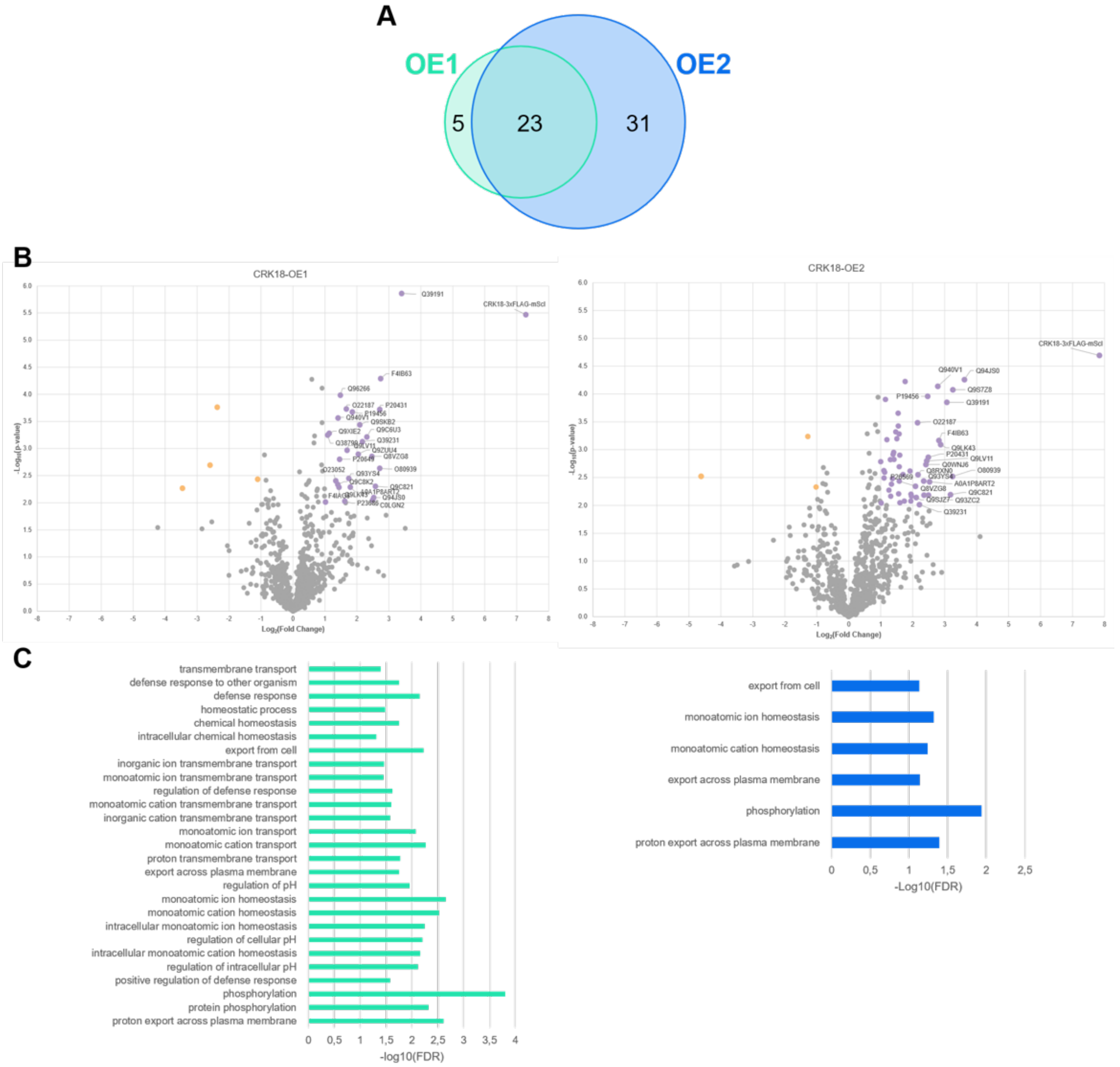


**Supplementary Figure 12. IP-MS of transgenic Arabidopsis CRK18 OE lines.** A) Comparison of proteins associated with CRK18 in CRK18-OE1 and CRK18-OE2. The Venn diagram of proteins significantly enriched (p-value < 0.01, log2(FC)> 1) by IP-MS of CRK18 in CRK18-OE1 and CRK18-OE2 in four-week-old Arabidopsis rosettes. B) Volcano plots of proteins identified by IP-MS in CRK18-OE1 and CRK18-OE2 compared to the control. Proteins which were significantly increased (p-value <0.01, log2FC >1) or decreased (p-value <0.01, log2FC >-1) in CRK18-OE1 or CRK18-OE2 compared to the control are marked in purple and orange, respectively. C) Gene Ontology (GO)-term analysis of proteins significantly enriched in CRK18-OE1 (left) and CRK18-OE2 (right).

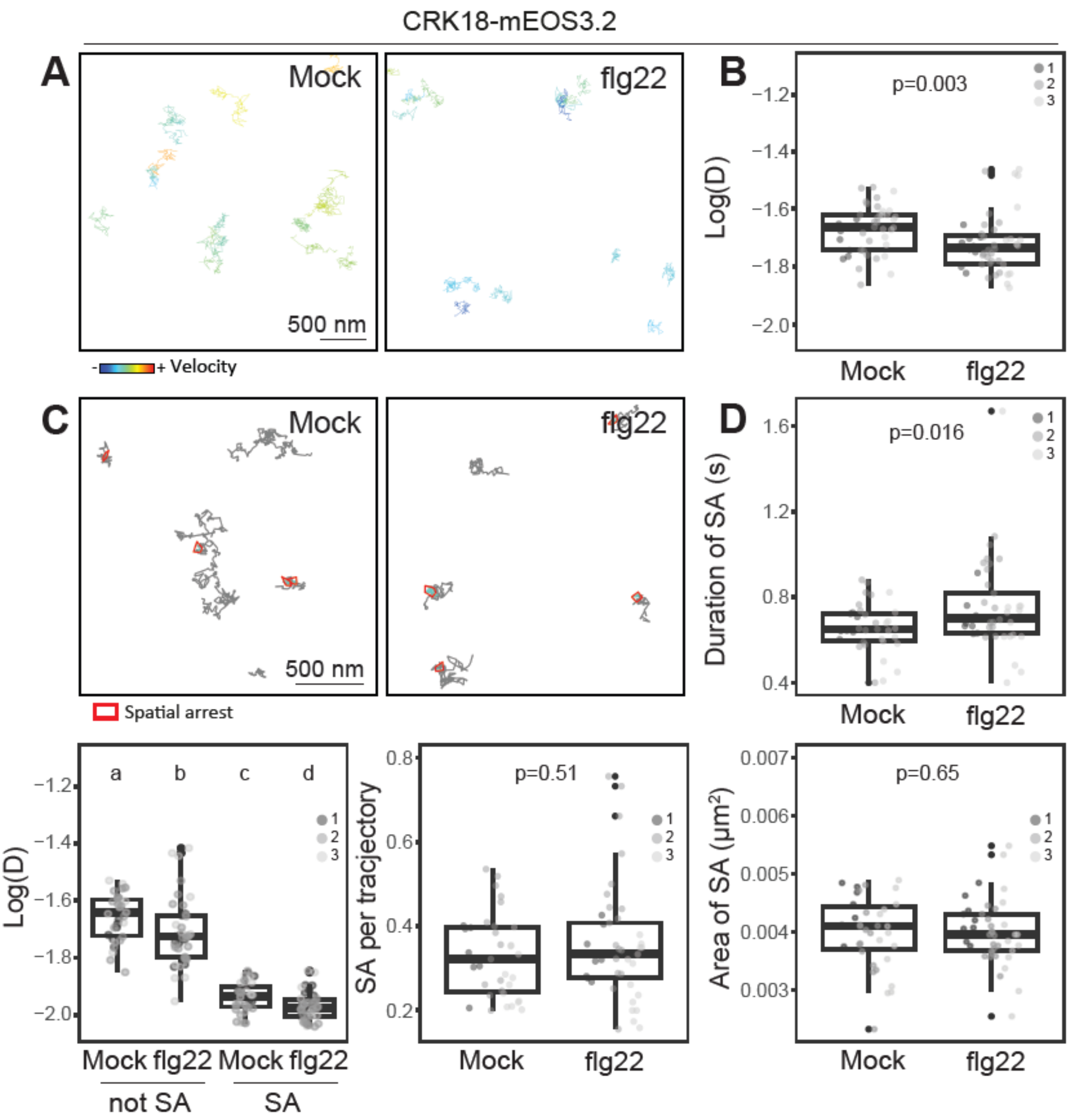


**Supplementary Figure 13. Long-term single-molecule imaging of CRK18-mEOS3.2.** A) Representative images of CRK18-mEOS3.2 long-term single molecule trajectories observed during transient expression in *N. benthamiana*, in presence of 1 µM flg22 or corresponding mock control ($H_2O$). Colours indicate single molecule velocity. B) Quantification of instantaneous diffusion coefficient (D) within 10 min of treatment. Each data point represents the average log(D) value obtained per cell (n), colours indicate independent experiments. C) Representative images of CRK18-mEOS3.2 single molecule spatial arrest (SA). D) Analysis of CRK18-mEOS3.2 spatial arrests (SA). Each data point represents the average duration of SA in seconds (s) (top right), the diffusion coefficient of not in spatial arrest (non-SA) and in spatial arrest (SA) molecules (bottom left), the number of spatial arrests per trajectory (bottom middle) and the average area of spatial arrest (bottom right) obtained per cell (n), colours indicate independent experiments. P values indicate statistical testing using pairwise Wilcoxon with Bonferroni correction. Conditions that do not share a letter are significantly different in pairwise Wilcoxon with Bonferroni correction ($p<0.05$). In total, n=35 (mock) and n= 41 (flg22) cells from three independent experiments were imaged and analysed.

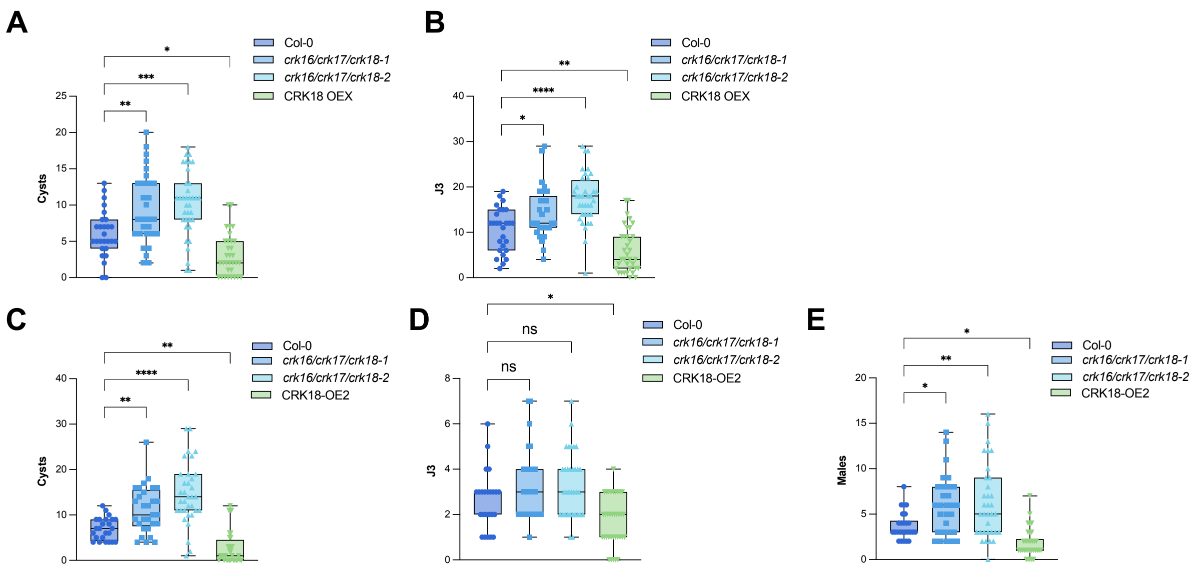


**Supplementary Figure 14. CRK18 modulate plant susceptibility to the cyst nematode *Heterodera schachtii*.** Two-week-old Arabidopsis seedlings were inoculated with ~250 H. schachtii juveniles. A-B) The number of infections was counted 14 days post-inoculation. The graph shows the number of cysts and J3 (developing juveniles). C-F) The number of infections was counted 28 days post-inoculation. The graph shows the number of cysts, J3 (developing juveniles) and males. Each experiment was performed three times with similar results (n=25-36), which were pooled and analyzed using One-way ANOVA followed by Dunn's post-hoc test with Benjamini-Hochberg correction for multiple comparisons (n=25-36).

**Supplementary Table 1. Effect of the disulphide bonds on the stability and localization of CRK18.** For each cysteine residue in the CRK18-ECD AF model, the solvent accessibility was calculated as a percentage of the available surface area. The position of the disulphide bonds in the AlphaFold structure of CRK18-ECDs are marked in orange. Localization of full-length CRK18 mutant variants in *N. benthamiana* and difference in Tm (ΔTm) of the CRK18 mutants (ΔTm = Tm (CRK18) - Tm (CRK mutant)) of the cysteine to alanine mutant disulphide pairs. Tm-values were derived from the thermal unfolding curves obtained at 350 nm, pH 5. We did not perform the thermal stability measurements for the other mutant variants, which were marked with ND (not determined).

| Cysteine | C30 | C106 | C82 | C91 | C94 | C122 | C204 | C213 | C216 | C241 | C227 | C228 |
|---|---|---|---|---|---|---|---|---|---|---|---|---|
| Solvent accessibility (%) | 34 | 0 | 8 | 3 | 11 | 0 | 0 | 0 | 0 | 0 | 63 | 1 |
| Position | 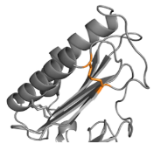 | | 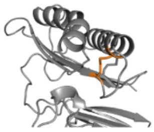 | | 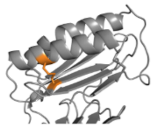 | | 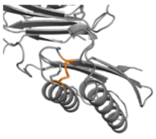 | | 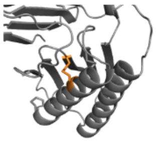 | | 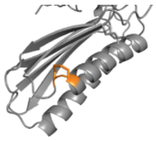 | |
| Cysteine mutant | C30A,C106A | | C82A,C91A | | C94A,C122A | | C204A,C213A | | C216A,C241A | | C227A,C228A | |
| $\Delta T_m$ (°C) | ND | | ND | | 12 | | ND | | 7 | | 6 | |
| Localization | ER | | ER | | ER | | ER | | ER | | PM | |

**Supplementary Table 2. Pairwise backbone RMSD (Å) between the representative coarse-grained structures selected for backmapping.** RMSD values (Å) calculated between the representative coarse-grained conformations selected from each independent CG simulation and subsequently used as starting structures for backmapping and all-atom molecular dynamics simulations.

| | Backmapped Replica 1 | Backmapped Replica 2 | Backmapped Replica 3 |
|---|---|---|---|
| **Backmapped Replica 1** | 0.00 | 1.33 Å | 1.70 Å |
| **Backmapped Replica 2** | 1.33 Å | 0.00 | 2.10 Å |
| **Backmapped Replica 3** | 1.70 Å | 2.10 Å | 0.00 |

**Supplementary Table 3. Structural and energetic comparison of the representative closed and open CRK18 ECD conformations.** Interface geometry, interaction energetics and binding affinity estimates calculated using PDBe PISA, PRODIGY and PRODIGY Crystal. These analyses identify the closed conformation as the biologically relevant dimeric assembly, whereas the more open conformation displays features consistent with partial interface destabilization.

| | Parameter | Closed conformation | Open conformation |
|---|---|---|---|
| **PDBe PISA Assembly** | Oligomeric state | Homodimer | Non-stable |
| | Buried interface area (Å$^2$) | 2254.2 | 1106.8 |
| | $\Delta G_{int}$ (kcal mol$^{-1}$) | −9.4 | - |
| | $\Delta G_{diss}$ (kcal mol$^{-1}$) | 0.9 | - |
| | Complexation Significance Score (CSS) | 1.00 | 0.00 |
| **PDBe PISA Monomer interface** | Interface residues (Chain A) | 35 | 20 |
| | Interface residues (Chain B) | 38 | 21 |
| | Buried surface (Chain A, Å$^2$) | 1155.9 | 569.6 |
| | Buried surface (Chain B, Å$^2$) | 1098.3 | 544.4 |
| **PRODIGY Protein-protein** | Binding free energy, ΔG (kcal mol$^{-1}$) | −11.3 | -8.9 |
| | Predicted $K_d$ (25 °C) | 4.9 nM | 310 nM |
| **PRODIGY Crystal** | Biological interface probability | 0.608 | 0.192 |
| | Crystallographic interface probability | 0.392 | 0.808 |
| | Link density | 0.070 | 0.110 |
| | Charged-charged contacts | 1 | 2 |
| | Charged-polar contacts | 7 | 3 |
| | Charged-apolar contacts | 1 | 7 |
| | Polar-polar contacts | 15 | 2 |
| | Polar-apolar contacts | 29 | 7 |
| | Apolar-apolar contacts | 17 | 12 |

**Supplementary Table 4. Predicted pKa values of CRK18 cysteine residues calculated with PROPKA for the representative closed monomer and homodimer conformations**. The representative closed conformation obtained from AA molecular dynamics simulations was used as input for PROPKA analysis. The table lists the predicted pKa values, PROPKA solvent accessibility classification, and the principal local determinants of the predicted pKa for each cysteine residue in both monomers of the homodimer. Chain A was extracted and used as the representative monomer for comparison.

| Cysteine | Monomer pKa | Dimer chain A pKa | Dimer chain B pKa | PROPKA environment | Major local determinants of predicted pKa |
|---|---|---|---|---|---|
| C30 | 9.36 | 11.71 | 13.78 | Buried | D118 |
| C82 | 12.89 | 9.81 | 9.97 | Buried | C94, C122 |
| C91 | 9.07 | 9.07 | 8.42 | Surface | R88 |
| C94 | 8.93 | 15.57 | 8.53 | Buried | E118, S116, C82, C122 |
| C106 | 9.14 | 14.51 | 10.11 | Buried | E118, R47 |
| C122 | 11.42 | 12.00 | 15.98 | Buried | T98, C82, C94 |
| C204 | 13.43 | 13.43 | 12.87 | Buried | C213, C216 |
| C213 | 8.52 | 8.52 | 9.68 | Buried | Y184 |
| C216 | 8.92 | 9.12 | 9.19 | Buried | R238 |
| **C227** | **7.10** | **7.22** | **6.72** | **Surface/ Buried*** | **C227** |
| **C228** | **5.31** | **5.31** | **6.13** | **Surface** | **C228, R229** |
| C241 | 13.97 | 14.08 | 12.06 | Buried | N220, C204, C216 |

* PROPKA classified C227 in chain A of the homodimer as buried despite its minimal burial score (0.05).

**Supplementary Table 5. Extended list of proteins identified in CRK18 overexpression lines by IP-MS**. IP-MS experiments were performed on two Arabidopsis transgenic lines, CRK18-OE1 and OE2. Proteins significantly enriched (-log10 p-value >1.3, log2FC>1) in OE1 or OE2 compared to control are shown. (Table continues on next page).

| Uniprot ID | TAIR ID | Gene Name | OE-1<br>Log2 FC | OE-1<br>-log10 p-value | OE-2<br>Log2 FC | OE-2<br>-log10 p-value | Protein Description | Cellular Localization |
|---|---|---|---|---|---|---|---|---|
| Receptor Kinases | | | | | | | | |
| Q8RX80 | AT4G23260 | CRK18 | 7,28 | 5,47 | 7,84 | 4,69 | Cysteine-rich receptor-like protein kinase 18<br>Involved in defence and cell death responses | Plasma membrane |
| Q39191 | AT1G21250 | WAK1 | 3,39 | 5,86 | 3,07 | 3,85 | Cell wall-associated receptor kinase 1<br>Polygalacturonic acid binding (pectin), EGF-Like - Ca binding domain, interacts with GRP3 (glycine rich protein 3) | Plasma membrane |
| F4IB63 | AT1G51805 | SIF3 | 2,74 | 4,29 | 2,82 | 3,17 | Stress induced factor<br>Leucine-rich repeat protein kinase family protein<br>Putative tandem malectin-like LRR domain | Plasma membrane |
| O80939 | AT2G37710 | LecRK-IV.1 | 2,71 | 2,63 | 3,25 | 2,52 | L-type lectin-domain containing receptor kinase IV.1<br>Legume (L)-type lectin | Plasma membrane |
| Q9C821 | AT1G52290 | PERK15 | 2,58 | 2,30 | 3,18 | 2,19 | Proline-rich extensin-like receptor-like protein kinase 15<br>Extracellular extensin-like flexible proline-rich region<br>PERKs are proposed as sensors of cell wall integrity or stress | Plasma membrane |
| C0LGN2 | AT3G14840 | LIK1 | 2,49 | 2,05 | | | LYSM-RLK1 interacting kinase 1 (LIK1)<br>Probable leucine-rich repeat receptor-like serine/threonine-protein kinase, LRR single malectin-like domain, involved in plant immunty | Plasma membrane |
| Q8VZG8 | AT4G08850 | MIK2 | 2,46 | 2,86 | 2,49 | 2,19 | MDIS1-interacting receptor-like kinase 2<br>Leucine rich repeat receptor kinase, forms a complex with serine rich endogenous peptides (SCOOPs) and BAK1 and elicits immune responses | Plasma membrane |
| Q9SKB2 | AT2G31880 | SOBIR1 | 2,09 | 3,44 | 1,58 | 2,43 | Supressor of BIR1, Leucine-rich repeat receptor-like serine/threonine/tyrosine-protein kinase<br>Involved in defence responses and cell death, negative regulator of floral abscission, complex with RLP23 and BAK1, necrosis and ethylene-inducing peptide 1-like proteins (NLPs) | Plasma membrane |
| O22187 | AT2G23200 | LETUM1 | 1,66 | 3,73 | 2,15 | 3,49 | RLK with a tandem malectin-like ECD, CrRLK1L family member<br>Involved in autoimmune responses | Plasma membrane |
| Q9LK43 | AT3G23750 | TMK4/BARK1 | 1,43 | 2,28 | 2,87 | 3,09 | Receptor-like kinase TMK4<br>Auxin signal transduction, phosphorylation of TAA1 | Plasma membrane |
| Q9FID9 | AT5G38990 | MEDOS1/<br>LETUM2 | | | 1,97 | 2,16 | RLK with a tandem malectin-like domain, CrRLK1Ls family member, growth adaptation upon exposure to metal Ions | Plasma membrane |
| Pores/channels/pumps | | | | | | | | |
| P20649 | AT2G18960 | AHA1 | 1,45 | 2,80 | 1,70 | 2,55 | ATPase 1<br>Proton export across plasma membrane | Plasma membrane |
| P19456 | AT4G30190 | AHA2 | 1,85 | 3,68 | 2,47 | 3,96 | ATPase 2<br>Proton export across plasma membrane | Plasma membrane |
| P20431 | AT5G57350 | AHA3 | 2,71 | 3,72 | 2,49 | 2,86 | ATPase 3<br>Proton export across plasma membrane | Plasma membrane |
| Q9LV11 | AT5G62670 | AHA11 | 1,69 | 2,97 | 2,44 | 2,80 | ATPase 11<br>Proton export across plasma membrane | Plasma membrane |
| Q39231 | AT1G22710 | SUC2 | 2,17 | 3,13 | 2,21 | 2,01 | Sucrose transport protein SUC2<br>Transports sucrose into the cell together with protons | Plasma membrane |
| Q93YS4 | AT5G06530 | ABCG22 | 1,74 | 2,45 | 2,34 | 2,44 | ABC transporter G family member 22<br>Water deprivation, transpiration | Plasma membrane |
| Q9C8K2 | AT1G51500 | ABCG12 | 1,40 | 2,34 | 1,92 | 2,62 | ABC transporter G family member 12<br>Export of fatty acids | Plasma membrane |
| Q9XIE2 | AT1G59870 | ABCG36 | 1,13 | 3,28 | 1,39 | 2,92 | ABC transporter G family member 36<br>Auxin efflux transporter | Plasma membrane |
| Q8RXN0 | AT1G17840 | ABCG11 | | | 2,16 | 2,55 | ABC transporter G family member 11<br>Export of fatty acids | Plasma membrane |
| Q8LPK2 | AT4G25960 | ABCB2 | | | 1,60 | 2,90 | ABC (ATP-binding casette) transporter B family member 2 | Plasma membrane |
| Q9S7Z8 | AT1G70940 | PIN3 | | | 3,26 | 4,08 | Pin-formed3, Auxin efflux carrier component 3<br>Auxin Efflux, binds IAA | Plasma membrane |
| Q9C8G5 | AT1G30360 | ERD4 | | | 1,37 | 2,48 | Early-responsive to dehydration 4<br>CSC1-like protein, Calcium activated cation channel | Plasma membrane |
| Protein folding/export/reuptake | | | | | | | | |
| Q9C6U3 | AT3G08030 | | 2,31 | 3,21 | | | Uncharacterized protein T8G24.2<br>DUF642 cell wall protein | Secreted |
| Q0WNJ6 | AT3G11130 | CHC1 | | | 2,41 | 2,74 | Clathrin heavy chain 1<br>Vesicle formation, endocytosis | Clathrin complex |
| Q9ZVJ6 | AT2G38750 | ANN4 | | | 1,55 | 3,43 | Annexin D4<br>Ca dependent membrane binding, Golgi mediated secretion, probably involved in aba signalling and osmotic stress | Golgi apparatus<br>Plasmodesmata |
| P23686 | AT1G02500 | SAM1 | 1,62 | 2,03 | | | S-adenosylmethionine synthase 1<br>S-adenosylmethionine catalysis | Cytosol |
| Q93ZC2 | AT5G36230 | | | | 2,35 | 2,19 | ARM repeat superfamily protein | Cytoplasm |
| Q94AW8 | AT3G44110 | ATJ3 | | | 1,41 | 2,95 | Chaperone protein dnaJ 3<br>Heat shock response, protein folding, iron binding | Cytoplasm |
| P42825 | AT5G22060 | ATJ2 | | | 1,31 | 2,82 | Chaperone protein dnaJ 2<br>Heat shock response, protein folding, iron binding | Cytoplasm |
| Q9SJ66 | AT2G35840 | SPP2 | | | 1,19 | 3,18 | Probable sucrose-phosphatase 2<br>Sucrose synthesis | Cytoplasm |

| Uniprot ID | TAIR ID | Gene Name | OE-1<br>Log2 FC | OE-1<br>-log10 p-value | OE-2<br>Log2 FC | OE-2<br>-log10 p-value | Protein Description | Cellular Localization |
|---|---|---|---|---|---|---|---|---|
| Chloroplastic | | | | | | | | |
| Q96266 | AT2G47730 | GSTF8 | 1,48 | 3,98 | 1,58 | 3,28 | Glutathione S-transferase F8<br>Toxin catabolism | Chloroplast |
| Q940V1 | AT4G39960 | DJA5 | 1,41 | 3,56 | 2,78 | 4,14 | Molecular chaperone Hsp40/DnaJ family protein<br>Heat shock response, protein folding,iron binding | Chloroplast |
| F4IAG5 | AT1G11750 | CLPP6 | 1,01 | 2,01 | | | ATP-dependent Clp protease proteolytic subunit<br>Proteolysis | Chloroplast |
| Q9SJZ7 | AT2G22360 | DJA6 | | | 2,11 | 2,14 | Chaperone protein dnaJ A6<br>Heat shock response, protein folding,iron binding | Chloroplast |
| Q9AST9 | AT1G73110 | | | | 1,94 | 2,08 | Ribulose bisphosphate carboxylase/oxygenase (RuBisCO) activase<br>P-loop containing nucleoside triphosphate hydrolases superfamily protein | Chloroplast |
| P54887 | AT2G39800 | P5CSA | | | 1,77 | 4,22 | Delta-1-pyrroline-5-carboxylate synthase A<br>Proline biosynthesis | Chloroplast,<br>Cytoplasm |
| Q9LIB2 | AT3G29320 | PHS1 | | | 1,60 | 2,05 | Alpha-glucan phosphorylase 1<br>Carbohydrate metabolism | Chloroplast |
| P46416 | AT5G27380 | GSHB | | | 1,58 | 2,69 | Glutathione synthetase<br>Glutathione biosynthesis | Chloroplast |
| Q8LPS6 | AT1G02150 | PPR3/CCR16 | | | 1,56 | 2,21 | Pentatricopeptide repeat-containing protein<br>Probable scaffolding protein | Chloroplast |

| | | | | | | | | |
|---|---|---|---|---|---|---|---|---|
| P17745 | AT4G20360 | EFTU/ATRAB8D | | | 1,55 | 3,66 | Elongation factor Tu<br>Facilitates RNA bidning to ribosomes | Chloroplast |
| Q9S713 | AT1G68830 | STN7 | | | 1,51 | 3,20 | Serine/threonine-protein kinase<br>Regulation of photosynthesis | Chloroplast |
| Q9SH69 | AT1G64190 | PGD1 | | | 1,47 | 3,32 | 6-phosphogluconate dehydrogenase<br>Oxidative decarboxylation, metabolism | Chloroplast |
| Q8H112 | AT4G22890 | PGL1A | | | 1,42 | 2,82 | PGR5-like protein 1A<br>Electron transport in photosystem I | Chloroplast |
| Q39102 | AT1G50250 | FTSH1 | | | 1,34 | 2,38 | ATP-dependent zinc metalloprotease FTSH 1<br>Repair of photosystem II | Chloroplast |
| Q9LIK9 | AT3G22890 | APS1 | | | 1,32 | 2,16 | ATP sulfurylase 1<br>Involved in sulfate assimilation pathway | Chloroplast |
| Q9LMR1 | AT1G15730 | ZNG2A2 | | | 1,26 | 2,27 | Cobalamin biosynthesis CobW-like protein<br>Similar to other ZNG1 metallochaperones | Chloroplast |
| Q9C9I7 | AT1G71500 | PSB33 | | | 1,15 | 3,90 | Photosystem B protein 33<br>Provides stability to Photosystem II | Chloroplast |
| Q9FH02 | AT5G42270 | FTSH5 | | | 1,13 | 2,59 | ATP-dependent zinc metalloprotease FTSH 5<br>Repair of photosystem II | Chloroplast |
| Q9SA52 | AT1G09340 | CP41B | | | 1,10 | 2,62 | Chloroplast stem-loop binding protein<br>Putatative RNA-binding protein | Chloroplast |
| O80860 | AT2G30950 | FTSH2 | | | 1,00 | 2,78 | ATP-dependent zinc metalloprotease FTSH 2<br>Repair of photosystem II | Chloroplast |
| Q9ZUU4 | AT2G37220 | CP29B | 2,04 | 2,90 | 1,95 | 2,20 | RNA-binding protein | Chloroplast |
| Q0WN54 | AT1G80030 | DJA7 | 1,79 | 2,29 | 2,52 | 2,43 | Molecular chaperone Hsp40/DnaJ family protein<br>Heat shock response, protein folding,iron binding | Chloroplast |
| Other | | | | | | | | |
| Q38799 | AT5G50850 | PDH2/MAB1 | 1,08 | 3,25 | 1,02 | 2,04 | Pyruvate dehydrogenase E1 component subunit beta-1<br>Part of the pyruvate dehydrogenase complex, convert<br>pyruvate to s acetyl-CoA and CO2 | Mitochondrion |
| Q94JS0 | AT5G13430 | UCR1-1 | 2,52 | 2,10 | 3,61 | 4,26 | Cytochrome b-c1 complex subunit Rieske-1<br>Iron sulfur cluster containing protein, part of a complex<br>that is involved in electron transport in the mitochondria,<br>important for oxidative phosphorylation to produce ATP | Mitochondrion |
| Q9ZT91 | AT4G02930 | TUFA | | | 1,11 | 2,49 | Elongation factor Tu<br>Likely promotes binding of tRNA to Ribosomes | Mitochondrion |
| Q570B4 | AT2G26250 | FDH | | | 1,74 | 2,08 | 3-ketoacyl-CoA synthase 10 (KCS-10), FIDDLEHEAD<br>Synthesis of lipids, Cuticular wax biosynthesis | Endoplasmic reticulum |
| P26569 | AT2G30620 | H12 | | | 2,09 | 2,35 | Histone H1.2<br>Nucleosome assembly, DNA binding | Nucleus |
| O23052 | AT1G05150 | | 1,33 | 2,41 | | | Calcium-binding tetratricopeptide family protein | ND |

**Supplementary Table 6. List of proteins differentially expressed the four-week-old rosettes of CRK18-OE1 and CRK18-OE2.** The table shows the differentially expressed proteins (FDR adjusted p-value < 0.05) for the two Arabidopsis CRK18 overexpression lines (OE1 and OE2) found in the total proteome compared to the control line. (Table continues on next page).

| **Upregulated** | | | **OE1** | | **OE2** | | | |
|---|---|---|---|---|---|---|---|---|
| **Uniprot ID** | **TAIR ID** | **Gene Name** | **$log_2$(FC)** | **p-value** | **$log_2$(FC)** | **p-value** | **Protein Description** | **Cellular Localization** |
| Q93Z66 | AT1G10700 | PRS3 | | | 0,44 | 0,035 | Phosphoribosyl pyrophosphate (PRPP) synthase3 | Chloroplast |
| Q84WN0 | AT4G37920 | CDB1 | 0,59 | 0,03 | | | Chloroplast development and biogenesis 1<br>Involved in chloroplast development | Chloroplast |
| Q8RX87 | AT5G20250 | RS6/DIN10 | | | 0,84 | 0,036 | Dark inducible 10, raffinose synthase 6<br>Glycosyl hydrolase | Cytoplasm |
| Q9C505 | AT1G69410 | ELF5A-3 | 0,53 | 0,002 | 0,45 | 0,002 | Eukaryotic elongation factor 5A-3<br>Putatative translation initation factor | Cytoplasm |
| Q93VP3 | AT1G26630 | ELF5A-2 | | | 0,38 | 0,029 | Eukaryotic elongation factor 5A-2<br>Translation initiation factor, involved in programmed cell death responses | Cytoplasm |
| P42734 | AT4G39330 | CAD9 | | | 0,48 | 0,020 | Probable cinnamyl alcohol dehydrogenase 9<br>Involved in monolignol biosynthesis | Cytosol |
| Q9FK25 | AT5G54160 | OMT1 | | | 0,33 | 0,043 | O-methyltransferase 1<br>Flavonol biosynthesis | Cytosol |
| Q9SSK5 | AT1G70890 | MLP43 | | | 0,39 | 0,024 | Major latex protein like 43<br>Uncharacterized | Cytosol, mitochondrion |
| P49211 | AT4G18100 | EL32Z | 0,62 | 0,01 | | | Large ribosomal subunit protein eL32z<br>Involved in translation | Cytosolic ribosome |
| Q9SR37 | AT3G09260 | BGLU23 | | | 0,62 | 0,022 | Beta-glucosidase 23<br>Component of ER bodies | Endoplasmic reticulum |
| Q9SE50 | AT1G52400 | BGLU18 | | | 0,43 | 0,042 | Beta-glucosidase 18<br>Inducible ER body formation, abscisic acid metabolism | Endoplasmic reticulum |
| Q9FF98 | AT5G23820 | ML3 | | | 0,62 | 0,044 | MD-2-related lipid-recognition protein 3<br>Involved in defence responses | Endoplasmic reticulum, vacuole |
| Q9FYH7 | AT1G30900 | VSR6 | 1,04 | 0,0001 | | | Vacuolar-sorting receptor 6<br>Protein targeting to vacuole | Golgi transport complex |
| Q940S0 | AT1G14670 | TMN2 | 0,49 | 0,03 | | | Transmembrane 9 superfamily member 2<br>Endomembrane protein | Golgi, membrane |
| Q8RXN4 | AT4G31810 | MS47 | 0,79 | 0,004 | | | Small ribosomal subunit protein mS47<br>Proteolytic activity | Mitochondrion |
| Q8RXW8 | AT5G65207 | | | | 0,78 | 0,019 | Uncharacterized protein | ND |
| O04658 | AT5G27120 | NOP5-1 | 0,63 | 0,02 | | | Probable nucleolar protein 5-1<br>Similar to SAR DNA-binding protein-1 | Nucleolus |
| P98204 | AT5G04930 | ALA1 | 0,33 | 0,03 | | | Aminophospholipid ATPase1<br>Putative aminophospholipid translocase | Plasma membrane |

| **Downregulated** | | | **OE1** | | **OE2** | | | |
|---|---|---|---|---|---|---|---|---|
| **Uniprot ID** | **TAIR ID** | **Gene Name** | **$log_2$(FC)** | **p-value** | **$log_2$(FC)** | **p-value** | **Protein Description** | **Cellular Localization** |
| Q9SW21 | AT4G25050 | ACP4 | | | -3,27 | 0,018 | Acyl carrier protein 4<br>Fatty acid biosynthesis | Chloroplast |
| Q9XI93 | AT1G13930 | | | | -1,29 | 0,016 | Nodulin-related protein 2<br>Involved in salt stress response | Chloroplast |
| Q9SD66 | AT3G47070 | | | | -0,72 | 0,017 | Thylakoid soluble phosphoprotein | Chloroplast |
| Q9XFT3 | AT4G21280 | PSBQ1 | -0,47 | 0,005 | | | Photosystem II subunit Q1<br>Part of oxygen-evolving complex of photosystem II | Chloroplast |
| P56777 | ATCG00680 | PSBB | -0,55 | 0,0001 | | | Photosystem II CP47 reaction center protein<br>Component of photosystem II | Chloroplast |
| Q682S0 | AT2G30520 | RPT2 | -0,40 | 0,03 | -0,49 | 0,00004 | Root phototropism protein 2<br>Phototropism response | Cytoplasm |
| Q9M7Q7 | AT1G07370 | PCNA1 | | | -0,57 | 0,001 | Proliferating cellular nuclear antigen 1<br>Putatatively involved in cell cycle regulation | Cytoplasm |
| P35510 | AT2G37040 | PAL1 | | | -0,60 | 0,001 | Phenylalanine ammonia-lyase 1<br>Cinnamic acid biosynthesis | Cytoplasm |
| P30187 | AT2G41090 | CML10 | | | -0,76 | 0,024 | Calmodulin-like protein 10<br>Calcium binding | Cytoplasm |
| Q8RXZ8 | AT3G09980 | ACIP1 | -0,41 | 0,03 | | | Acetylated interacting protein1<br>Microtubule associated, bacterial immunity | Cytosol |
| P93017 | AT2G33830 | DRM2 | -0,84 | 0,04 | | | Dormancy-associated gene 2<br>Negative regulator of immunity | Cytosol |
| Q9ZQ80 | AT2G03440 | NRP1 | -0,96 | 0,0001 | -0,76 | 0,001 | Nodulin-related protein 1<br>Involved in stress responses, potential regulator of ABA | Cytosol |
| Q9LJ86 | AT3G22600 | LTPG5 | | | -1,27 | 0,023 | Glycosylphosphatidylinositol (GPI)-anchored lipid protein transfer 5<br>Lipid transport, PAMP signalling | Cytosol, extracellular region, membrane |
| F4HQT8 | AT1G13470 | | | | -0,82 | 0,044 | Hypothetical protein, insecticidal crystal toxin domain-containing protein | Extracellular region |
| Q208N7 | AT1G06460 | ACD321 | -0,52 | 0,04 | | | Alpha-crystallin domain 32.1<br>Homology to small heat shock proteins | Extracellular region, peroxisome |
| Q94BV7 | AT4G05020 | NDB2 | | | -0,52 | 0,007 | NAD(P)H dehydrogenase B2<br>Mitochondrial alternative NADH dehydrogenase | Mitochondrion |
| O49447 | AT4G28390 | AAC3 | | | -0,72 | 0,032 | ADP-ATP carrier protein 3<br>Antiporter involved in ADP/ATP transmembrane transport | Mitochondrion |
| A0A1P8B0G6 | AT2G38890 | | | | -2,94 | 0,028 | Uncharacterized protein | ND |
| O80644 | AT2G39570 | ACR9 | -2,60 | 0,01 | | | ACT domain-containing protein ACR9<br>Amino acid binding, metabolic pathways | Nucleus |

| | | | | | | | | |
|---|---|---|---|---|---|---|---|---|
| O80644 | AT2G39570 | ACR9 | -2,60 | 0,01 | | | ACT domain-containing protein ACR9<br>Amino acid binding, metabolic pathways | Nucleus |
| P43286 | AT3G53420 | PIP2A | -0,73 | 0,02 | | | Plasma membrane intrinsic protein 2A<br>Aaquaporin, water transport | Plasma membrane |
| F4K518 | AT5G61910 | | | | -1,51 | 0,0002 | DCD (Development and Cell Death) domain protein<br>Calcium binding, cell response | Plasma membrane |
| Q9SRQ6 | AT3G03520 | NPC3 | -0,80 | 0,0001 | | | Non-specific phospholipase C3<br>Lipid synthesis | Vacuole, plasma membrane |

**Supplementary Table 7. Extended data of phosphoproteomics of CRK18-OE1 and CRK18-OE2 Arabidopsis lines.** The table shows the significant (FDR-adjusted p-value < 0.05) hits for each overexpression line (OE1 and OE2) compared to the Control.

| Upregulated | | | | OE-1 | | OE-2 | | | |
|---|---|---|---|---|---|---|---|---|---|
| Uniprot ID | TAIR ID | Gene Name | Phosphosite | log2 FC | p-value | log2 FC | p-value | Protein Description | Cellular Localization |
| Q9LE80 | AT3G15450 | | S218 | 2,8 | 0,02 | 3,3 | 0,00 | Aluminum induced protein with YGL and LRDR motifs | Chloroplast |
| Q9LFA4 | AT3G52870 | | S408 | | | 2,8 | 0,03 | IQ calmodulin-binding motif family protein | Chloroplast |
| Q6NMC1 | At1g72640 | | S84 | | | 2,0 | 0,02 | NAD(P)-binding Rossmann-fold superfamily protein | Chloroplast |
| Q96500 | AT3G26740 | LIR1,CCL | S114 | | | 1,7 | 0,01 | Light-regulated protein 1<br>Circadian controlled mRNA levels | Chloroplast |
| O80860 | AT2G30950 | FTSH2 | S393 | | | 1,2 | 0,02 | ATP-dependent zinc metalloprotease FTSH 2<br>Repair of photosystem II | Chloroplast |
| Q9ZTZ7 | AT1G01790 | KEA1 | S120 | | | 1,0 | 0,02 | K(+) efflux antiporter 1<br>Involved in chloroplastic $K^+$ homeostasis | Chloroplast |
| P42737 | AT5G14740 | BCA2 | S265 | | | 1,3 | 0,02 | Beta carbonic anhydrase 2<br>Carbon utilization | Cytoplasm |
| Q9LIL3 | AT3G22850 | | S215 | 1,1 | 0,03 | | | Aluminum induced protein with YGL and LRDR motifs | Cytosol, plasma membrane |
| O64650 | AT2G45710 | RPS27A | S80 | 1,3 | 0,05 | | | Small ribosomal subunit protein eS27z<br>Zinc-binding ribosomal protein | Cytosolic ribosome |
| Q0WV86 | AT3G48200 | | S808 | | | 1,7 | 0,03 | Transmembrane protein | Membrane |
| Q8RX80 | AT4G23260 | CRK18 | T498 | | | 3,8 | 0,02 | Cysteine-rich receptor-like protein kinase 18<br>Involved in defence and cell death responses | Plasma Membrane |
| Q93XY1 | At5g53620 | | S341 | | | 1,7 | 0,02 | RNA polymerase II degradation factor | Plastid |
| Q0WMN5 | At3g49140 | HUGZ-2 | S355 | | | 1,6 | 0,02 | HugZ domain containing protein<br>Part of the CLPPRT complex network | Plastid |

| Downregulated | | | | OE-1 | | OE-2 | | | |
|---|---|---|---|---|---|---|---|---|---|
| Uniprot ID | TAIR ID | Gene Name | Phosphosite | log2 FC | p-value | log2 FC | p-value | Protein Description | Cellular localization |
| Q8RWQ4 | AT5G19390 | PHGAP2 | S753 | -1,4 | 0,03 | -1,4 | 0,02 | Rho GTPase-activating protein 7<br>Activator for Rac-type GTPase | Cytoplasm, Chloroplast,<br>Plasma Membrane |
| F4JKH6 | AT4G28080 | REC2 | S1293 | | | -1,1 | 0,04 | Protein REDUCED CHLOROPLAST COVERAGE 2<br>Involved in chloroplast development | Cytosol |
| F4JAE0 | AT3G53390 | | S107 | -1,4 | 0,04 | | | Transducin/WD40 repeat-like superfamily protein | Nucleus |
| F4ISN0 | AT2G25730 | | S1698 | | | -1,2 | 0,04 | Zinc finger FYVE domain protein | Nucleus |
| Q9XER9 | AT5G23150 | HUA2 | S185 | -4,0 | 0,02 | | | ENHANCER OF AG-4 protein 2<br>Putative transcription factor | Nucleus |
| Q39032 | AT5G58670 | PLC1 | S260 | -1,2 | 0,02 | -1,0 | 0,05 | Phosphoinositide phospholipase C 1<br>Induced by environmental stresses, Involved in production of second messengers | Plasma Membrane |
| Q8W4J9 | AT5G43470 | RPP8 | S142 | -1,3 | 0,02 | | | Recognition of Peronospora parasitica 8<br>Resistance to Peronospora parasitica | Plasma Membrane |
